# Computational simulations reveal that Abl activity controls cohesiveness of actin networks in growth cones

**DOI:** 10.1101/2021.11.01.466771

**Authors:** Aravind Chandrasekaran, Akanni Clarke, Philip McQueen, Hsiao Yu Fang, Garegin A. Papoian, Edward Giniger

## Abstract

Extensive studies of growing axons have revealed many individual components and protein interactions that guide neuronal morphogenesis. Despite this, however, we lack any clear picture of the emergent mechanism by which this nanometer-scale biochemistry generates the multi-micron scale morphology and cell biology of axon growth and guidance *in vivo*. To address this, we studied the downstream effects of the Abl signaling pathway using a computer simulation software (MEDYAN) that accounts for mechanochemical dynamics of active polymers. Previous studies implicate two Abl effectors, Arp2/3 and Enabled, in Abl-dependent axon guidance decisions. We now find that Abl alters actin architecture primarily by activating Arp2/3, while Enabled plays a more limited role. Our simulations show that simulations mimicking modest levels of Abl activity bear striking similarity to actin profiles obtained experimentally from live-imaging of actin in wild type axons in vivo. Using a graph-theoretical filament-filament contact analysis, moreover, we find that networks mimicking hyperactivity of Abl (enhanced Arp2/3) are fragmented into smaller domains of actin that interact weakly with each other, consistent with the pattern of actin fragmentation observed upon Abl overexpression *in vivo*. Two perturbative simulations further confirm that high Arp2/3 actin networks are mechanically disconnected and fail to mount a cohesive response to perturbation. Taken together, these data provide a molecular-level picture of how the large-scale organization of the axonal cytoskeleton arises from the biophysics of actin networks.

**Highlight summary:** How do single-molecule dynamics produce multi-micron scale changes in actin organization in an extending axon? Comparison of computational simulations to in vivo data suggests that Abl kinase and Arp2/3 expand actomyosin networks by fragmenting into multiple domains, thus toggling the axon between states of local vs global internal connectivity.

## Introduction

During embryonic development, neurons send out axonal projections to construct complex neural circuits. (Araújo and Tear, 2003) The ability of axons to extend through the extracellular space *in vivo* is coordinated by specialized structures found at the tips of motile axons, called growth cones, that span tens of microns. (Dent *et al*., 2011) Protrusive forces are generated in the growth cone through complex biochemical interactions between membrane-bound signaling molecules that intercept extracellular guidance cues (Netrins, Semaphorins, Slit, and Ephrin) (Huber *et al*., 2003; Hinck, 2004; Seiradake *et al*., 2016; Feinstein and Ramkhelawon, 2017) and the intracellular cascades that transmit these signals to force producing cellular components, namely the cellular cytoskeleton. Thus, growth cones are made of actin and microtubule filaments along with a host of actin modifying proteins and signaling molecules such as Rho-family GTPases.(Giniger, 2002; Huber *et al*., 2003; Gallo and Letourneau, 2004) Actin and microtubules play specific roles in the extension of axons, where actomyosin drives motility and guidance, while microtubules are thought to provide structural support and lock in the consequences of actin dynamics. (Dent and Gertler, 2003; van Goor *et al*., 2012; Kahn and Baas, 2016; Turney *et al*., 2016) Many studies have tried to dissect the mechanism of axon growth by focusing on the morphological changes of growth cones or the cytoskeletal signaling processes that aid in growth. (Witte and Bradke, 2008) Despite detailed understanding of the various of components involved in the process, however, we lack any clear picture of an emergent mechanism that can explain how the biochemically identified nanometer level changes to signaling molecules and cytoskeletal binding proteins lead to micron level changes to axonal architecture.

Studies focusing on growth cone morphology have investigated the relationship between structural dynamics of the cytoskeleton and the mechanism of axon extension. Fluorescent and electron microscopic imaging of neuronal cell cultures has revealed two major modes of axonal growth (Strasser *et al*., 2004; Sánchez-Soriano *et al*., 2010; Tamariz and Varela-Echavarría, 2015). Cultured neurons often display axons with flat, fan-shaped growth cones (Forscher and Smith, 1988; Schaefer *et al*., 2002; Gallo and Letourneau, 2004; Lowery and Vactor, 2009; Suter and Forscher, 2010) However, growing evidence from *in vivo* studies primarily reveals growth cones that are bulbous and only minimally adherent to the substratum, extending by the formation and selective stabilization of filopodial protrusions.(O’Connor *et al*., 1990; Grabham *et al*., 2003; Omotade *et al*., 2017; Clarke *et al*., 2020a, 2020b; Santos *et al*., 2020) The cytoskeletal organization and dynamics of this *in vivo* class of growth cones has received far less attention in the past, and it is the relationship between signaling and actin organization in growth cones such as these on which we focus in the current study.

The growth cone architecture during axonal protrusion is controlled by signal transduction molecules that dynamically alter kinetic fluxes of actin-binding proteins to modify cytoskeletal structure. One critical signaling module studied extensively in axons is the Abelson (Abl) nonreceptor tyrosine kinase. Abl is known to act downstream of many of the widespread, phylogenetically conserved axon guidance receptors. (Jones *et al*., 2004) Genetic perturbations that alter Abl activity in *vivo* increase the probability of developmental errors due to axonal misrouting and stalling. Genetic and biochemical analyses have suggested that Abl alters growth cone organization by simultaneously stimulating activity of the branching actin nucleator, Arp2/3, while inhibiting the processive actin polymerase, Enabled (Ena) -. (Kannan and Giniger, 2017; Kannan *et al*., 2017) While the roles of these two critical downstream proteins on the actin cytoskeleton have been studied independently (Pollard, 2007; Winkelman *et al*., 2014; Harker *et al*., 2019b; Ni and Papoian, 2019), the relationship and relative roles of the two are still poorly understood. (Bear *et al*., 2002) Live *in vivo* imaging studies of an identified axon in the developing Drosophila wing, called TSM1, revealed that the baseline, wild-type level of Abl activity ensures spatial correlation and dynamic coherence of actin networks within the growth cone. Under both Abl overexpression and Abl knockdown conditions, growth cones are characterized by increased spatial fragmentation of actin and dynamic incoherence. Additionally, Abl regulates growth cone size, with Abl over-expressing axons having longer growth cones, while axons from neurons with reduced Abl have shorter growth cones.

Despite knowledge of the biochemical targets of Abl and its consequences at the growth cone level, we do not understand how these are linked through the dynamics of actin. As it is not possible to resolve single actin filaments by *in vivo* imaging experiments in live tissue, here we use a sophisticated computational model called MEDYAN to simulate key cytoskeletal processes that result from Abl signaling and then compare those simulations to the effects of Abl we have quantified previously *in vivo*. As biochemical and genetic experiments have identified Arp2/3 and Ena as crucial targets of Abl, we simulate networks with 20µM actin and a wide range of Arp2/3 and Ena concentrations. Critically, we perform these simulations at spatial and temporal scales that are large enough to capture properties of the TSM1 growth cone. We find that Arp2/3 plays a crucial role in determining actin filament length distributions and actin packing in growth cones. We also see that the growth cone simulations capture essential features of actin distributions measured in live-imaging experiments and reveal evidence to explain the mechanism of actin fragmentation observed in Abl-overexpressing axons *in vivo*. Finally, we study the functional consequences of the actin architectures formed in response to varying levels of Arp2/3 and Ena and show that Abl overexpression causes mechano-chemical disconnection within the actin network. We discuss the potential consequences of fragmentation for the motility, growth, and guidance of axons such as TSM1.

## Results

In order to faithfully simulate axonal actin networks, we simulated contractile actin networks under cylindrical boundary conditions using MEDYAN (Popov *et al*., 2016), a powerful active matter simulation software. As simulations over the entire length of a growing axon are computationally expensive, we restricted the bulk of our simulations to cylindrical reaction volumes of length 7.5µm and radius 1μm. This volume represents approximately 50% of the growth cone volume observed *in vivo* for TSM1, a growth cone with uniquely well-characterized actin distribution and dynamics that we take as our point of comparison throughout this analysis. (Clarke *et al*., 2020a) Axonal growth cone mimics were generated through stochastic evolution of actin networks initialized with 400, 40 monomer long filaments that were randomly oriented in the reaction volume with 20μM actin. Actin filaments are restricted to the reaction volume through a reflective boundary that also slows down the polymerization rates of filament ends in accordance with the Brownian ratchet model (Mogilner and Oster, 1996). Mole ratios of non-muscle myosin minifilaments (NM-IIA) and α-actinin were chosen to be 0.1 and 0.01, respectively, to observe contractile behaviors similar to previous studies. (Floyd *et al*., 2019; Li *et al*., 2020) Please refer to Methods for more information on MEDYAN and Supplementary Tables S1 and S2 for parameters used in the simulations employed for this study.

To understand the effect of Abl on actin architecture, we chose to model key downstream effectors of Abl, namely Arp2/3 and Ena. In MEDYAN, branched nucleation is modeled as a chemical event where freely diffusing Arp2/3 complex binds to a parent filament to produce an offspring filament at approximately 70º angle. (Mullins *et al*., 1998; Pollard, 2007) In this study, we have also incorporated the force-sensitive unbinding of Arp2/3 nucleated branches.(Fujiwara *et al*., 2007; Pandit *et al*., 2020) The concentration limits for Arp2/3 and Ena were obtained using a minimalistic ordinary differential equation (ODE) model of actin network dynamics to identify the range over which network properties are sensitive to varying concentration. Please refer to Supplementary Methods in Supplementary Information for more details on the ODE model. To understand the effect of Abl on the actin cytoskeleton, we simulated networks at 1, 5, 10, 25, and 50nM of Arp2/3 and Ena each with six replicates of 2000s long trajectories and analyzed as shown below.

The rest of the Results section is organized as follows. We start by analyzing the actin filament properties that are observed in our simulations when varying the two downstream Abl effectors, Arp2/3 and Ena. We then quantify micron-scale actin network organization in the simulations using metrics that correspond to properties measured in previous experimental analyses (actin spread and actin pair-correlation profile) or that illuminate the basis of those properties (modularity of actin networks). We then investigate the functional significance of network organization and axonal coherence with two perturbative experiments. Finally, we compare our computational results with experimental observations in vivo in TSM1 axons.

### Filament branching versus extension – Arp2/3 driven changes dominate filament length

MEDYAN simulations reveal that Arp2/3 and Enabled change the length distribution of actin filaments, but with Arp2/3 playing the predominant role. Representative final snapshots shown in Figure 1 show marked differences in actin network organization that result from changing Arp2/3 and Ena concentrations. Please refer to Supplementary Information, Figures S1-S4 for enlarged images along each axis. We quantified filament length distributions to define the specific changes resulting from varying Arp2/3 and Ena (Figure 2A, corresponding probability density functions are shown in Supplementary Information, Figure S5). As Arp2/3 concentration is increased, Arp2/3 kinetics causes increased branched filament nucleation (Figure 2B). As the total actin in the reaction volume is held constant, this results in an increased abundance of short filaments. Changing Ena concentration, in contrast, has a limited impact on filament length distribution. As Ena concentration increases, we see a steady increase in the fraction of filament plus ends stabilized by Ena (Figure 2C). At [Arp2/3] <10nM, addition of Ena caused Ena-driven filament extension leading to increased abundance of longer filaments (>1μm) and decreased abundance of shorter filaments (<1μm). At [Arp2/3] ≥10nM, changing Ena concentration has a limited impact on filament lengths (Figure 2A and Supplementary Information, Figure S5).

**Figure 1.**
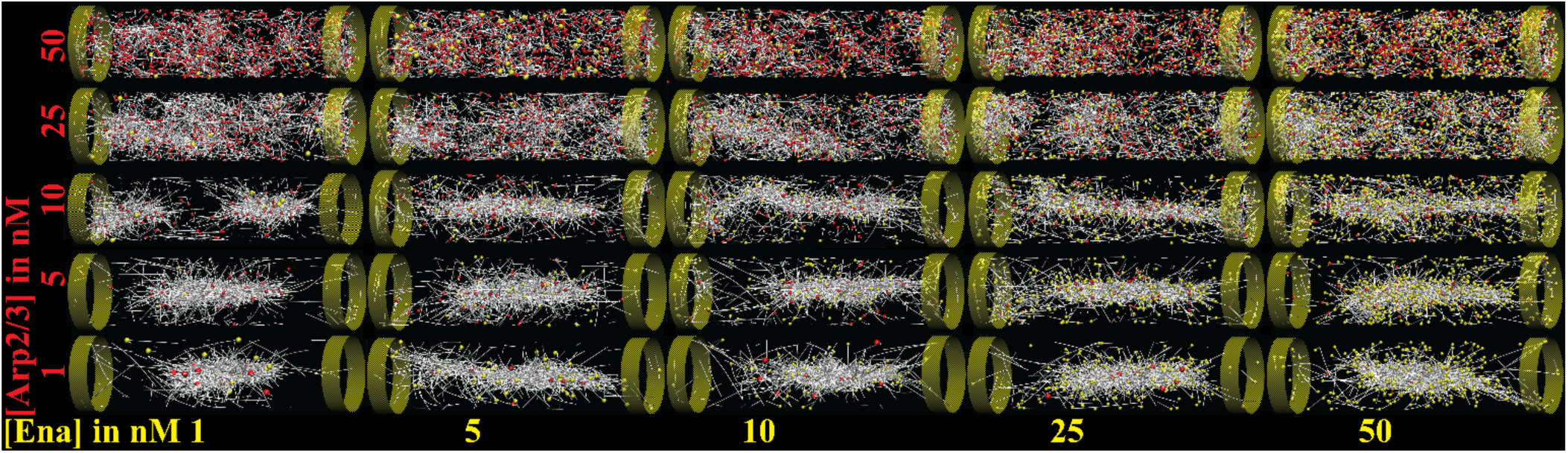
Gallery of representative, final snapshots from 2000s long simulations of 20μM actin in cylindrical volumes of length 7.5µm and radius 1µm at various Arp2/3 and Ena concentrations. Arp2/3 concentrations are mentioned at the bottom, while Ena concentrations are noted to the left. F-actin bound Arp2/3, and Ena molecules are colored in white, red, and yellow, respectively. Myosin minifilaments and α-actinin are also present in the simulations but are not visualized. Supplementary Information, Figures S1-S4 show enlarged images along each row and column.

**Figure 2.**
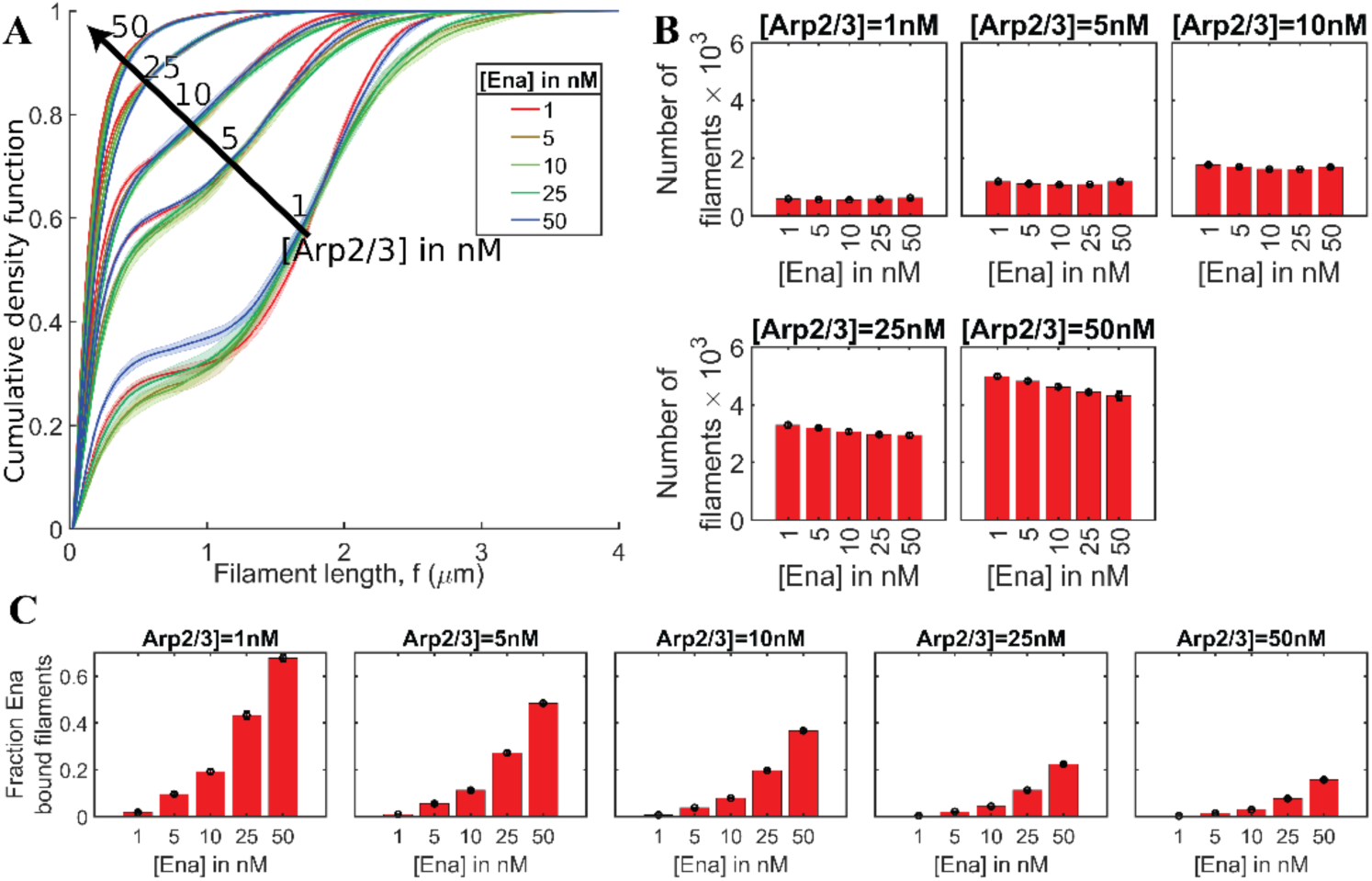
Arp2/3 dynamics determines filament organization of growth cone mimics. A. Filament length distributions are shown as cumulative density functions at various Arp2/3 and Ena concentrations. When calculating filament length distributions, parent filament and daughter filaments were considered as two different filaments. The corresponding probability density functions are shown in Supplementary Information, Figure S5. Distributions are colored based on Ena concentration, while the Arp2/3 concentration is overlaid on the graph. Solid lines and shaded areas represent mean and standard deviation, respectively, from multiple replicates. B. Bar graph shows the number of filaments when Ena concentrations are changed at each Arp2/3 concentration. Error bars show standard deviation. C. Bar graph shows the fraction of Ena bound filaments found at the end of the simulations. Error bars show standard deviation.

### Arp2/3 and Ena alter network organization at different length scales

We next found that Arp2/3 also dominates the overall organization of the total actin network. To understand how differences in filament lengths alter network organization, we quantified actin concentration in 100nm thick discs along the cylindrical reaction volume axis (as shown in Figure 3A schematic). Averaged 1-dimensional profiles of actin distribution along the volume were obtained by using convolution to peak-align the actin profiles from the last 200s of the trajectories (Figure 3B). Increasing Arp2/3 at any given Ena concentration leads to drastic changes in actin profiles characterized by multiple peaks spread across an increasing fraction of the reaction volume. In contrast, increasing Ena at low Arp2/3 concentrations modestly broadens the width of a central peak in the actin distribution, while at higher Arp2/3, increasing Ena has little discernable effect. To understand the relative effects of Arp2/3 and Ena on the actin distribution profiles, we measured the spread of actin around the mean (Figure 4). Actin spread was defined as the sum of the square root of second moments on either side of the distribution mean. (Clarke *et al*., 2020a) We then quantified the significance of Arp2/3 and Ena on actin spread using the Wilcoxon rank-sum test. We find that in 94% (47/50) of pairwise comparisons, increasing Arp2/3 concentration at any particular Ena concentration increases the median spread of actin profile at 0.05 significance. Similarly, at a given Arp2/3 concentration, increasing Ena increases actin spread in 86% (43/50) of comparisons. We note that at the two highest Arp2/3 concentrations, actin is already distributed roughly homogeneously across the entire volume, limiting the possibility of further spread. We also note that increasing Arp2/3 increases actin spread over a multi-micron length scale at any given Ena concentration, whereas the range of actin spread from varying Ena is smaller than 1µm in all cases. Thus, these results suggest that actin organization and distribution are dominated by Arp2/3, with modest effects due to changes in Ena.

**Figure 3.**
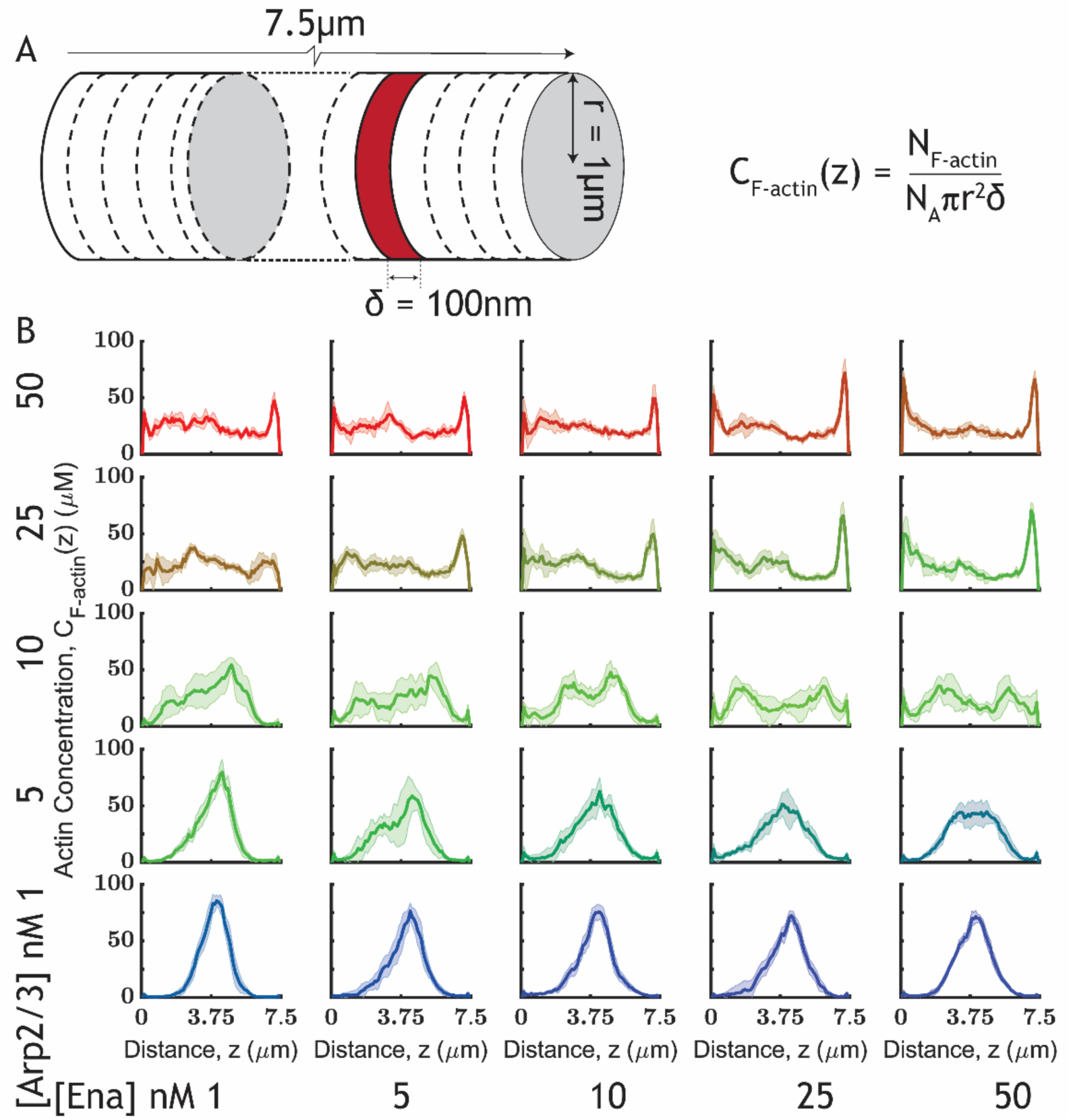
Axial actin concentration profile reveals Arp- and Ena-driven differences in actin organization. A. Schematic shows cylindrical reaction volume divided into discs of width 100nm. Local concentration of actin was defined based on the number of F-actin monomers in the disc at any given snapshot. B. Actin concentration in 100nm slices along the length of the reaction volume is plotted for the last 200s of trajectory calculated from all replicates at Arp2/3 and Ena concentrations. The solid line and shaded area represent mean and standard deviation, respectively. Arp2/3 concentrations are mentioned along Y-axis while Ena concentration is varied along X-axis.

**Figure 4.**
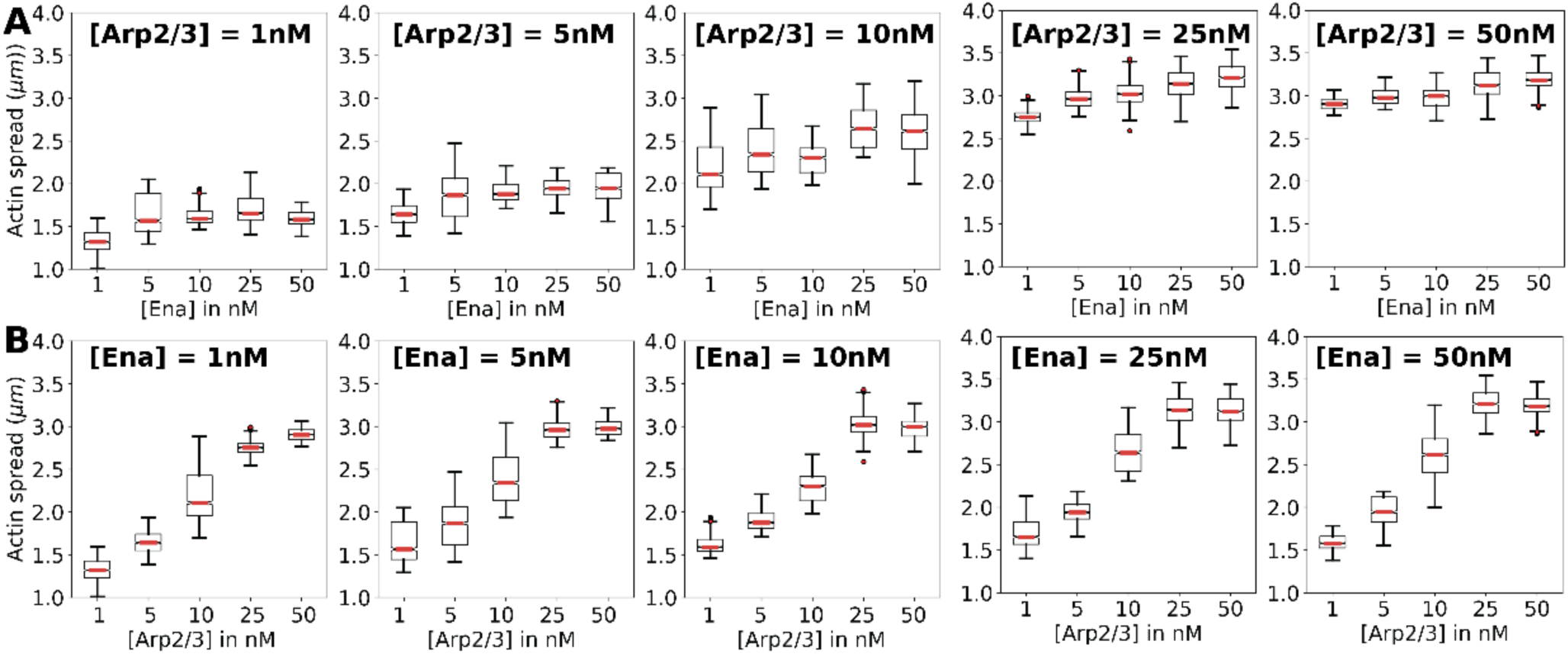
Actin spread from the last 200s of trajectories reveals Arp2/3 dependent actin organization in axon mimics. Actin spread was measured as the sum of left and right square root of the second moment from the actin distribution mean. A. Each sub-panel shows actin spread at various Ena concentrations at a given Arp2/3 concentration (mentioned above each panel). B. A. Each sub-panel shows actin spread at various Arp2/3 concentrations at a given Ena concentration (mentioned above each panel).

Further analysis was done to understand the role of the reaction volume dimensions in determining the actin distributions. As simulations in 15µm length scales are computationally expensive, we chose to simulate a subset of the Arp2/3 and Ena concentrations we have explored so far. As Abl signaling simultaneously promotes Arp2/3 activity and inhibits Ena activity (Kannan *et al*., 2017), we simulated actin networks at the conditions specified along the falling diagonal of Figure 1 inside reaction volumes that were 15µm long and 1µm in radius. We see that under conditions that mimic elevated Abl expression, i.e., under ([Arp2/3],[Ena]) pair values of (50,1), and (25,5) actin spreads homogeneously throughout both the short and long reaction volumes (Supplementary Information, Figure S6), though in the short volume we do observe some pile-up of actin at the ends of the interval. In contrast, in conditions with Arp2/3 and Ena concentration pair values of (10,10), (5,25), and (1,50), we see that actin is highly condensed at certain parts, evidenced by peaks in the axial actin distributions. Additionally, we also see that the width of actin peaks is similar in actin distributions from both 7.5 µm and 15 µm reaction volumes. Finally, the overall distribution of actin within 15 µm long reaction volumes under all conditions studied appears to approximate a concatenation of multiple copies of the actin profiles obtained at 7.5 µm length scales (albeit with some ‘filling-in’ of the interval between actin peaks at the highest Ena concentration in the large volume). These similarities between simulations from the two length scales support the argument that the actin distributions obtained at 7.5 µm length scales largely display features that arise from the biophysical properties of the network, not from the spatial limits of the simulation volume, and that are also observed in volumes with length scales comparable to experimental growth cones.

In addition to the overall longitudinal profile of the actin distribution, it was essential to obtain a quantitative measure of the internal organization of the actin network. In our previous experimental analysis of TSM1 actin organization in vivo, we used wavelet analysis of the 1-D actin profiles to determine the spatial distribution of actin within the mass of growth cone actin (Clarke *et al*., 2020b). We therefore performed a calculation that gives us an analogous relative measure of the internal distribution of actin density along the long axis of the simulation volume, called variously, the pair correlation profile or the radial distribution function of the actin profile. In brief, actin density is calculated in transverse slices at 50nm intervals along the length of the volume. These allow us to calculate the pairwise correlation of density as a function of distance away from any reference point in the reaction volume, giving us a relative measure of the spatial distribution of actin density along the long axis of the simulation volume when compared with bulk linear density (Please refer to Supplementary Information, Figure S7 for a schematic explaining the calculation) Thus, values above 1.0 indicate that actin abundance is above bulk density while values below 1.0 correspond to regions with actin below linear bulk density (moles of actin per nm). We performed this calculation for all the combinations of Arp2/3 and Ena concentrations studied. (Figure 5).

**Figure 5.**
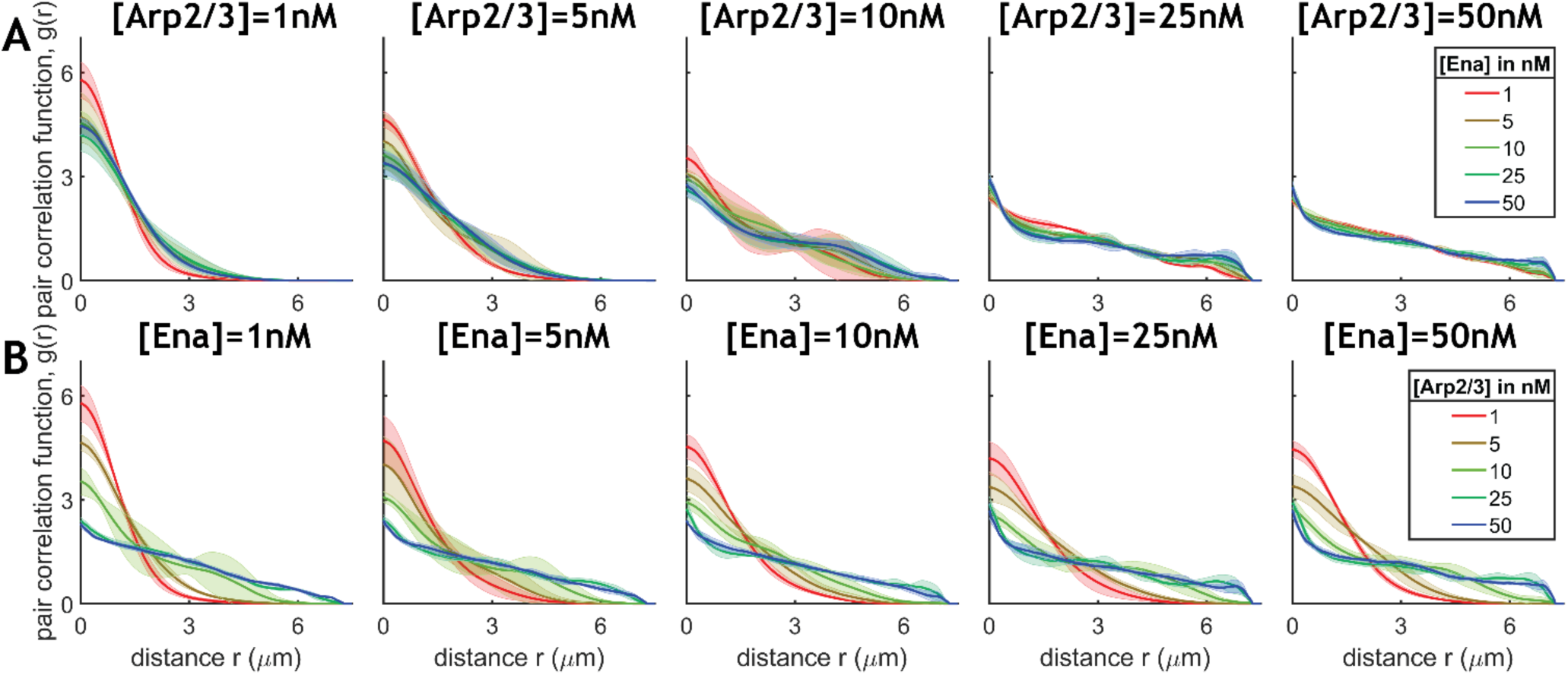
Pair correlation function of actin intensity reveals Arp dependant fragmentation. Mean (solid line) and standard deviation (shaded area) are plotted at various Ena (top row) and Arp2/3 (bottom row) concentrations at a given Arp2/3 and Ena concentration, respectively. These profiles were calculated by projecting monomer coordinates along the cylinder axis. Therefore, as two monomers can have the same coordinate along the cylinder axis, the radial pair correlation functions (Figure 7) are non-zero at 0µm. Please refer to Supplementary Information, Figure S11 for a schematic of the method used here.

We see that at Arp2/3 concentrations of 1nM and 5nM, the point-to-point correlation of F-actin density decreases monotonically with increasing distance. In other words, the slices of high actin density are clustered, with the probability of finding another high-density slice dropping monotonically with distance from the reference point along the longitudinal axis consistent with the single peak profiles shown in Figure 3B. Additionally, as Ena concentration is increased at these levels of Arp2/3, pair correlation decays to 1.0 (where local density falls to bulk density values) at slightly larger length scales. This result is consistent with the finding that actin spread increases modestly with increasing Ena at low Arp, as discussed earlier (compare Figure 3B and Figure 4A). At higher Arp2/3 concentrations, pair-correlation shows reduced dependence on distance, consistent with the relatively flat axial actin distribution profiles across the reaction volume (Figure 5). Changing Ena has very small effects on the pair correlation function profiles in these conditions, again consistent with actin spread data.

In contrast, when Arp2/3 concentration is increased at any given Ena concentration (Figure 5B), we see that linear actin densities are above bulk value (g(r)>1) over longer spans consistent with the increased actin spread observed earlier (Figure 4B). Specifically, the effect of Arp2/3 spans the entire reaction volume, consistent with the finding that Arp2/3 increases spread globally, across the entire volume, whereas Ena increases actin spread only locally, at short range. Finally, we see that at [Arp2/3]=10nM, the pair correlation function shows one auxiliary peak suggesting multiple domains of actin organization. At higher Arp2/3 concentrations, actin profiles decay monotonically at a slower rate, with no significant features. This is consistent with the complex and highly variable collection of peaks observed in the 1-D simulation profiles at high Arp2/3. The patterns in order parameters measured at high Arp2/3 concentrations along the cylinder axis, such as actin spread and pair correlation function, are mechanistically a consequence of actin fragmentation (explored in detail elsewhere; Chandrasekaran, et al., submitted). Thus, changing Arp2/3 concentration has a profound effect on the internal organization of the actin network, with low Arp2/3 promoting consolidation of actin in a homogeneous mass, and increasing Arp2/3 spreading actin into multiple, apparently separated masses. In the Discussion, we will analyze in detail the striking correspondence between these patterns of actin distribution in the simulations vs. those observed experimentally by imaging of actin in TSM1(Clarke *et al*., 2020b).

### Networks corresponding to elevated Abl activity are composed of multiple, weakly interacting actin filament communities

We wished to understand why increasing the level of the branching actin nucleator, Arp2/3, expands the actin network. Quantitative investigation of network connectivity now revealed that increasing Arp2/3 causes the actin to self-assemble into multiple, internally connected domains that are only weakly associated with one another. Modularity is a graph theoretical measure of networks to quantify whether and how network components are divided into modules, and such graph-theoretical metrics have proven helpful for defining network architecture. (Floyd *et al*., 2019; Eliaz *et al*., 2020). We constructed graph representations of inter-filament contacts by representing actin filaments as the nodes and weighting the edge by the total number of the linker, motor, and brancher contacts between any two filaments. We then compartmentalize the filaments into “communities” by optimizing the Louvain-Modularity metric. (Blondel *et al*., 2008) More details about the modularity metric can be found in Supplementary Information, Supplementary Methods. Communities identified by the algorithm are sets of actin filaments that interact more closely with other filaments within their own set than with the rest of the filaments in the network. We find that increasing Arp2/3 concentration increases the number of communities and decreases community size, suggesting that actin is organized into communities that only weakly interact. (Figure 6A and B). This altered organization of actin filaments also reduces the amount of actin in each cluster as Arp2/3 increases, as shown in Figure 6B.

**Figure 6.**
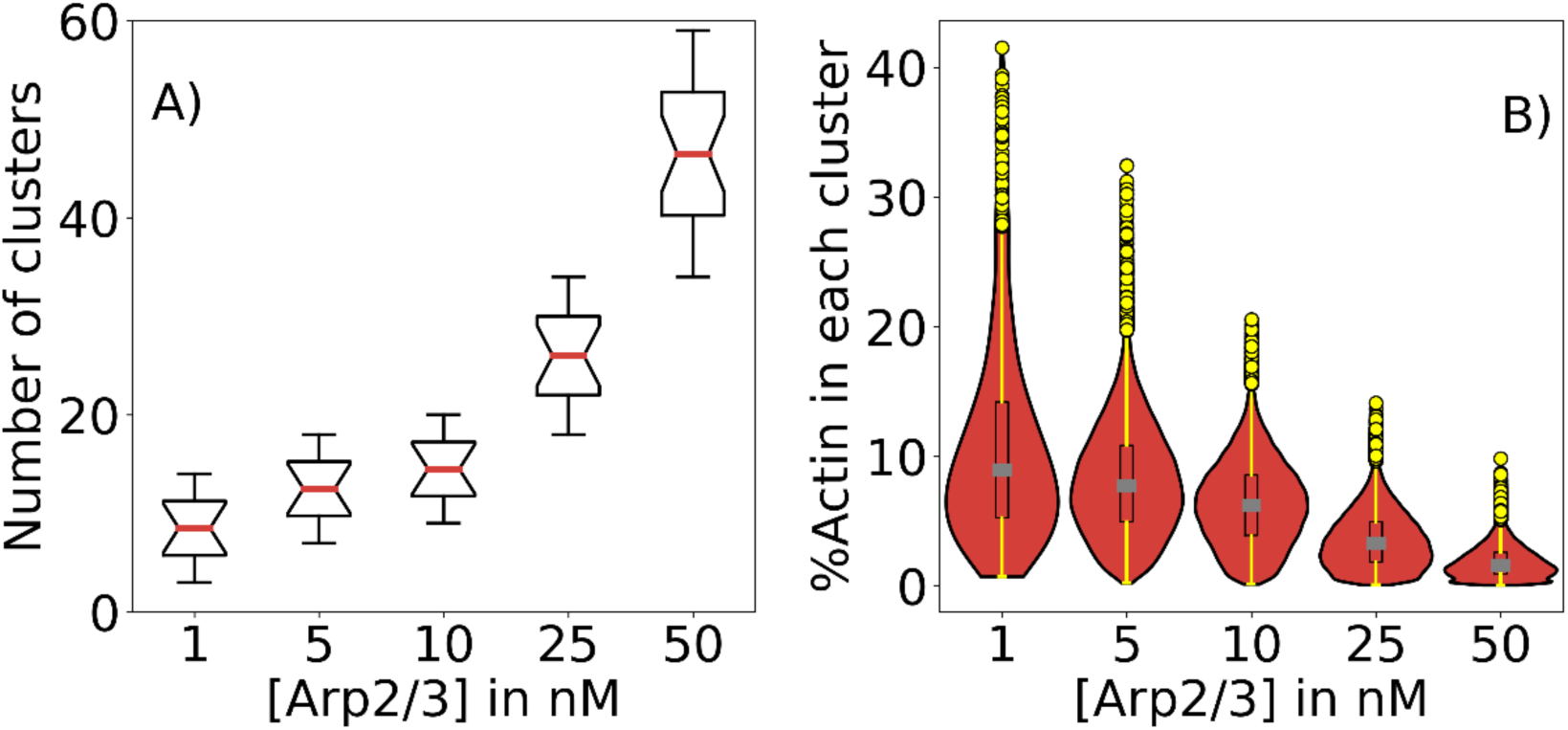
Arp2/3 activity alters network organization in favor of weakly coupled communities of filaments. A. The number of actin filament communities detected by the Louvain-Modularity optimization algorithm is plotted as a function of Arp2/3 concentration at [Ena] =25nM. Box and whisker plots show median (red), 25-75 percentile represented as black boxes, whiskers are shown as black lines, interquartile range (black whiskers) B. The fraction of F-actin in each cluster has been plotted as a violin plot. The box plot shows the 25-75 percentile represented as a box with the median shown as gray lines. Whiskers are shown as solid yellow lines with outlier data points represented as yellow circles

### Actin communities in elevated Arp2/3 networks are mechano-chemically disconnected

Functional analysis of mechanical connectivity and mechano-chemical response to perturbation reveal mechanical disconnection of actin networks as Arp2/3 level is increased. First, we asked if the weakly coupled communities that we find in high Arp2/3 networks are mechanically disconnected. We employed the final configurations of the 2000s long simulations presented above as the initial conditions for this perturbative study. To examine just the mechanical response of the system, we prevented chemical dynamics in filaments (polymerization, depolymerization, branching, and unbranching), thereby freezing any myosin and crosslinker molecules in the bound state. Localized myosin and crosslinker activity were then allowed only in a single 500nm “active zone” that was chosen to lie at the region of highest actin density. This localized activity was allowed to proceed for 500 seconds (full parameters of the simulation can be found in Supplementary information, Table S2). This simulation was inspired by previous experimental studies investigating spatial correlation in linear actin networks. (Linsmeier *et al*., 2016) Looking at the velocity of filaments from the last 100s of trajectories, we see that filaments move within the active zone and in a region adjacent to it (Figure 7). Specifically, in low Arp2/3 networks, actin filament movement extended in regions flanking the active zone over longer distances than filaments in high Arp2/3 networks. The difference in velocity decay rates between low Arp2/3 and high Arp2/3 networks demonstrates that the increasing number and decreasing size of filament-filament contact-based communities is associated with a decreased range of mechanical connection of a given zone of the actin network to the remainder of the network.

**Figure 7.**
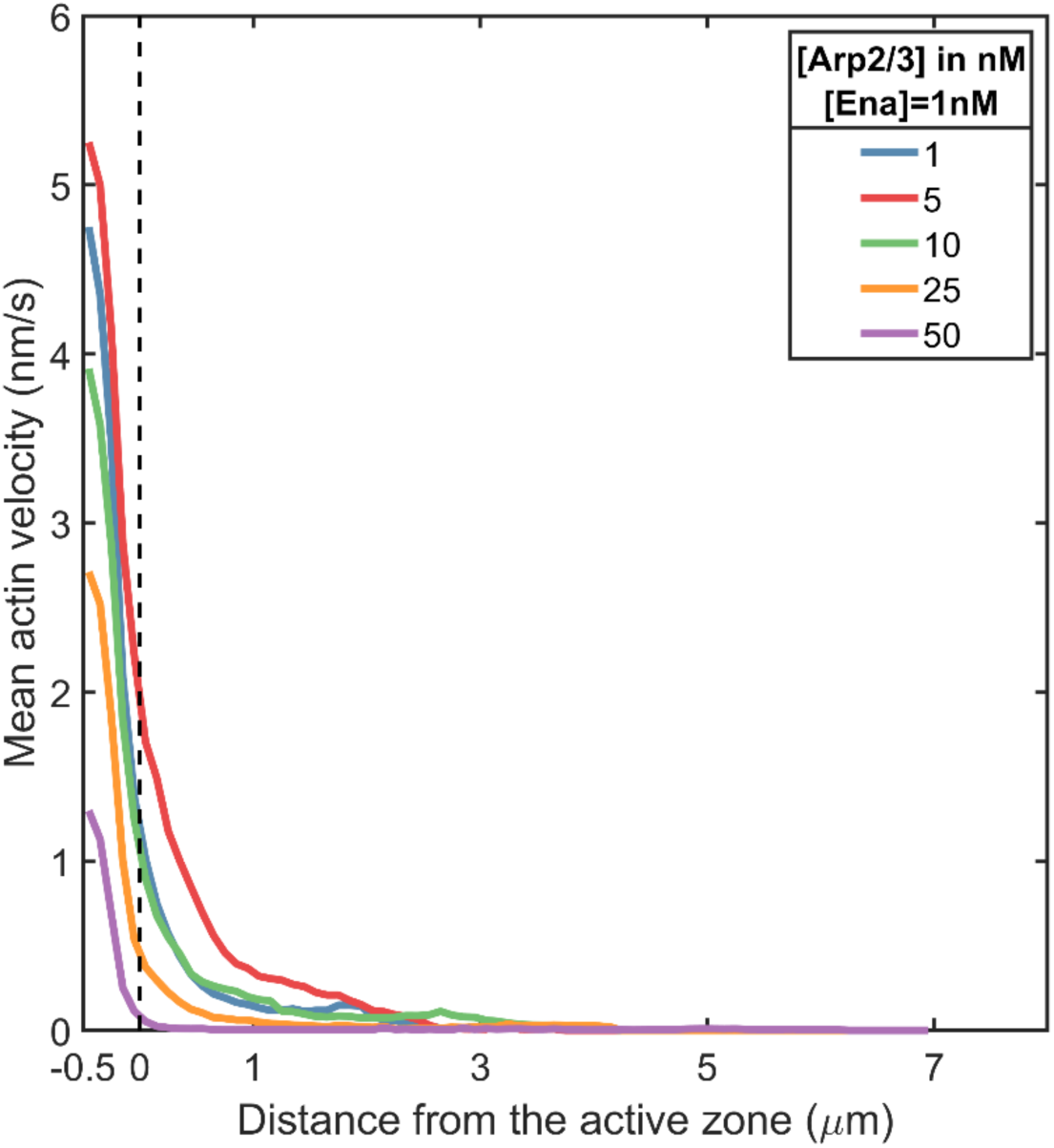
Mean filament velocity from spatially-localized myosin-driven perturbations reveals signs of mechanical fragmentation. Mean actin velocity calculated from displacements in the last 100s of trajectory at various Arp2/3 concentrations is shown. The dotted line represents the active zone boundary. N = 3 starting configurations for each Arp2/3 concentration indicated.

We next complemented this analysis of the structural connectivity of a static network by investigating the mechano-chemical response of an active actin network to mechanical perturbation. Again using the final configuration of filaments, linkers, motors, and branchers from our earlier simulations as the initial conditions, we exert an external force on the actin network using a mimic of a functionalized “Atomic Force Microscope (AFM) tip”. The AFM tip mimic is functionalized with 50 stabilized actin filaments of 540nm length (200 monomers). It is introduced at one end of the simulated growth cone network, close to a region of high actin density, and allowed to interact and become incorporated into that network through crosslinker and myosin driven mechanochemical couplings. With its associated stabilized actin filaments, the AFM tip mimic was then moved by a distance of 1µm over a 5.0s time scale. As a result, tensile forces are transmitted across the actin network through contacts between stabilized filaments in the probe and the actin filaments in the network. Chemical interactions such as filament treadmilling and inter-filament chemical interactions from branching, myosin, and crosslinker kinetics remained active throughout the reaction volume. We then visualize the response of the network to the force transmitted through the motions of the AFM probe. Networks are subject to three pull events and Figure 8 shows snapshots before and after each displacement of the AFM probe. In addition, we also show the “after” snapshot with each filament colored by the net displacement that it undergoes during the pull event. We observe stark differences in how actin networks at various Arp2/3 concentrations respond to this perturbation. We see that networks with [Arp2/3]=1nM and 5nM move coherently as a single connected unit in response to the external force. We also see that the AFM tip couples effectively with the rest of the network after the first pull event leading to significant filament displacements in the subsequent two pull events. On the other hand, in networks generated with [Arp2/3]=10nM and 25nM, AFM pulling reveals a functional disconnect between actin domains, preventing effective transmission of tension across the entire network. The displacement visualization also suggests that the very few filaments responded to the local probe displacement at these higher Arp2/3 concentrations. Taken together, the results from the perturbative simulations suggest that elevated Arp2/3 levels result in network architecture fragmented into domains that are mechano-chemically disconnected.

**Figure 8.**
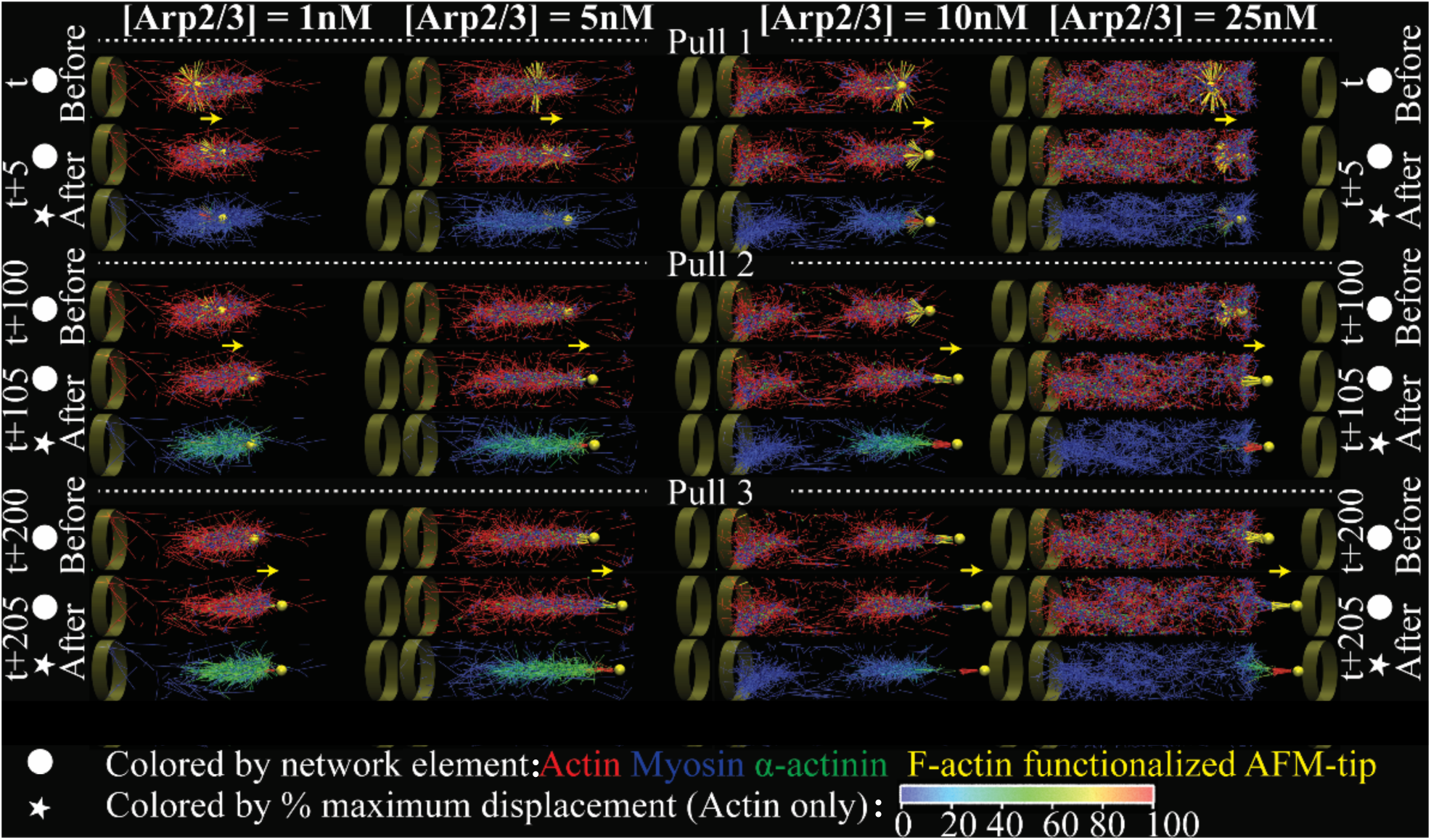
Mechanochemical response of actin networks to AFM mimic driven mechanical perturbations reveal fragmentation in intra-network signal transduction. Columns1-4 show time series of actin network snapshots (time stamp shown to the left and right) at various Arp2/3concentrations ([Ena]=1nM in all conditions shown) adapting to mechanical perturbation by a functionalized AFM probe (probe displacement velocity 200nm/s) in a cylinder of 9.5μm length and 1μm radius. Networks are subject to three AFM probe pull events (labelled Pull1, Pull2, and Pull3) each lasting 5s followed by regular network evolution for 95s. For each pull event, three snapshots are shown (one snapshot before and two after tip displacement). The snapshots before and after the AFM probe displacement show actin network (red) along with myosin (blue) and crosslinkers (green). The embedded AFM tip (yellow sphere) and the functionalized actin filaments attached to the tip (yellow cylinders) are also shown. The solid yellow arrow between the two snapshots represents the direction of displacement. Note that Arp2/3 and Ena molecules are present in the system but are not visualized. The third snapshot in each group visualizes actin filaments alone after the tip displacement, along with AFM tip, with filaments colored based on %maximum displacement (color bar shown below the image). Please note that the final snapshot only shows filaments that are present throughout the pull event. Snapshots corresponding to [Arp2/3]=50nM are shown in Supplementary Information, Figure S8, and video of the simulations shown above are included as Supplemental Movie M1.

### Actin organization within in silico networks resembles that of WT *Drosophila* axons

The simulations above were designed to model the properties of actomyosin networks. A key question, however, is how similar these are to actin distributions observed experimentally. We have previously used live fluorescent imaging to quantify the actin distribution in the growth cone of a growing axon in vivo in the developing Drosophila wing (TSM1 axon). To obtain representations of those experimental actin distributions comparable with the simulation results shown in Figure 3, we normalized the intensity profiles obtained from fluorescent imaging of actin in single WT axons over 30 minutes (sampling frequency = 3min), aligned the profiles by the position of maximum actin intensity in each time point and averaged the intensity for a 7.5µm span around that peak position. This procedure was repeated for 14 WT axons (presented as a gallery in Figure 9, showing the mean and standard deviation for each axon; Supplementary Information, Figure S9 shows actin profiles from MEDYAN simulations at all conditions studied alongside the experimental wild type profiles). Comparing the experimentally observed actin to the simulated profiles presented in Figure 3, we see that the experimental profiles show a distribution similar to that observed in the simulations at [Arp2/3] ≤10nM, with a dominant central peak surrounded by a steady decrease in actin intensity to either side.. Moreover, expanding the analysis of the experimental data to a total length of 15 μm reveals appearance of subsidiary peaks spaced a few microns from the central peak in many trajectories, and bears a marked resemblance to the additional peaks observed in simulated actin profiles in calculations performed in 15μm long cylinders with [Arp2/3] ≤10nM (Supplementary Information, Figure S10). Thus, even though our simulations approximate Abl signaling by just the downstream effector concentrations and do not include any neuronal polarization mechanisms, the actin profiles obtained from our simulations at [Arp] ≤ 10nM bear a striking similarity to the experimentally observed features of actin organization discussed above. This similarity is also evident in overlays of the individual profiles from the constituent experimental and simulation profiles (Supplementary Information, Figures S8). In contrast, it is clear from inspection of Figure 3 that simulations with higher levels of Arp2/3 do not bear any substantial resemblance to the experimentally observed axial actin distributions. Beyond the similarity in the overall profile of the experimental and simulated actin profiles, quantitative analysis reveals strikingly similar features in the internal organization of the actin density, and in the response of both profile and organization to perturbations of Abl-Arp2/3 signaling. These are summarized in Supplementary Information, Table S3, and will be considered in detail in the Discussion (below).

**Figure 9.**
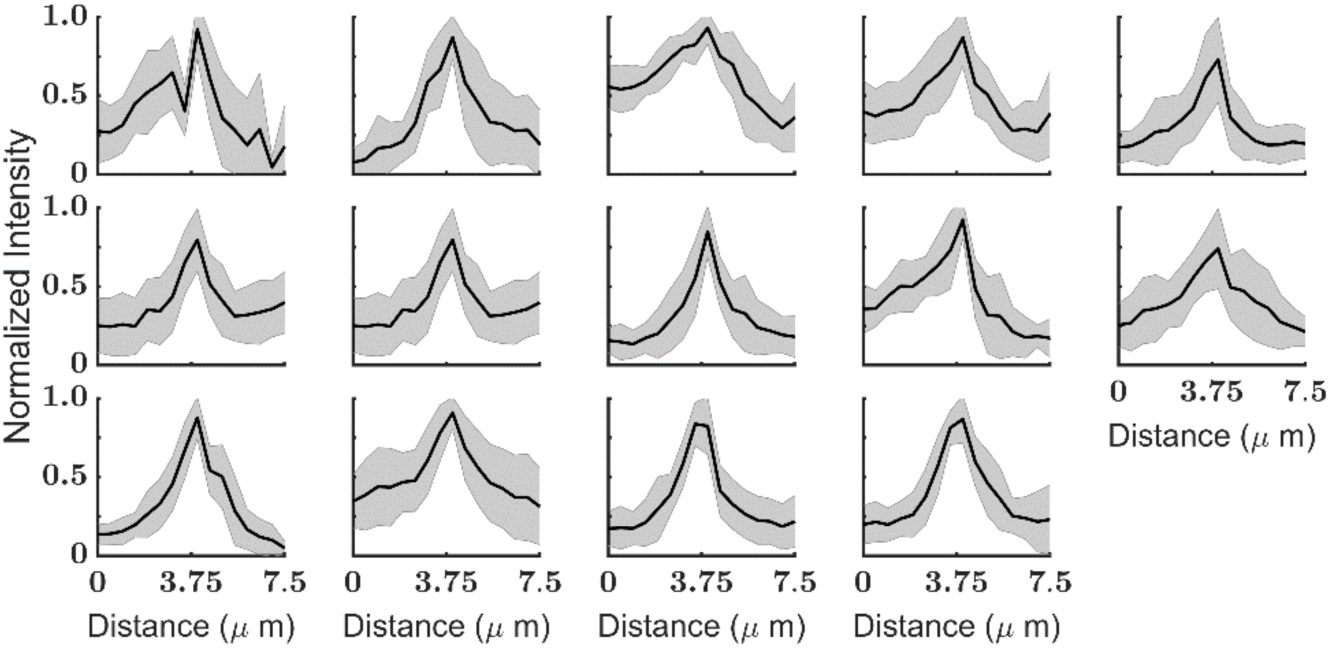
Simulations capture features of axonal growth cones observed in experiments. Actin intensity distribution in 14 WT Abl axons is shown in each sub-panel. The mean and standard deviation of peak-aligned actin profiles from the first 30 minutes of imaging a WT axon is plotted as a solid line and shaded area, respectively. Experimental actin intensity distributions were truncated 3.75 µm on either side of the peak. Note that the peak of experimental distributions was set to 3.75μm to aid comparison with the simulation data.

## Discussion

Axon growth and guidance are mediated by cytoskeletal dynamics that result from signaling cascades. In Drosophila, as in vertebrates, Abl is a crucial signaling molecule that controls axon growth by activating the actin nucleator, Arp2/3, and inhibiting the actin polymerase, Ena. Current methods, however, do not allow us to directly image the dynamics of individual actin filaments in growth cones that are embedded in living tissue. Therefore, to understand actin organization within an Abl-guided axon, we employed simulations that approximated the growth cone as a cylinder, varied Arp2/3 and Ena levels independently, and calculated their predicted effects on actin architecture. These showed that Arp2/3 and Ena affect actin organization at different length scales and magnitudes. Specifically, we found that Arp2/3 dynamics determine overall actin distribution, as it is a much more potent modulator of filament length and network organization than is Ena. While both Arp2/3 and Ena increase the spread of the actin network within the reaction volume, Ena locally broadens features of the actin distribution by tenths of a micron by extending individual actin filaments, whereas Arp2/3 globally spreads the actin distribution across multiple microns by fragmenting the actin into separate, weakly coupled domains. The functional consequences of this fragmentation are revealed by two perturbative simulations showing that actin networks become mechano-chemically decoupled under elevated Abl-Arp2/3 activity into separate domains of actin that fail to act cohesively. Finally, we show that our computational networks show a marked resemblance to the actin distributions observed in our previous quantitative imaging of the Drosophila TSM1 growth cone *in vivo*. Below, we present quantitative comparisons that further support the concordance between these simulations and previous live imaging data. We also discuss how the simulation results can account mechanistically for the actin distributions observed experimentally in both wildtype axons and axons with altered Abl activity, and why the potential for variable fragmentation and condensation of actin we demonstrate here may be essential to the normal dynamics of axonal actin in the wildtype context and how they may contribute to mutant phenotypes such as axon misrouting in the case of Abl-overexpressing axons.

As Abl steers the growth cone through changes to both Arp2/3 (promotion) and Ena (inhibition), how do we understand the relative role of each of the two downstream effector molecules on F-actin? The pronounced sensitivity of actin networks to changes in Arp2/3 over that of Ena is evident from our analysis of the spread and internal organization of the actin distributions. We define the spread of the distribution as the standard deviation of actin density about the midpoint of the volume. While increasing Ena at any given Arp2/3 concentration spreads the actin by no more than 0.1 – 0.3µm, changing Arp2/3 at any Ena spreads actin distribution on the order of several microns. Consistent results are obtained with profiles from 7.5 micron and 15 micron-long reaction volumes, suggesting that these observations are not due to actin-boundary interactions. The internal organization of the actin, in contrast, is a measure of how homogeneously actin is packed in the volume. By calculating the pairwise correlation of actin density as a function of distance along the long axis of the cylindrical volume, we can assess whether all of the actin is concentrated in a single, continuous domain or is split into local concentrations of actin separated by regions of reduced concentration. Pair correlation of actin along the cylindrical axis shows that Ena-driven spread has limited impact on domain organization, while Arp2/3 driven spread could arise from the separation of the underlying actin into discrete domains. An independent graph-theoretical modularity analysis confirmed these results suggesting fragmentation of actin due to Arp2/3 activity. Thus, we see that the network architecture reflects the presence of a variable number of domains, dependent on Arp2/3 activity and characterized by a combination of domain size and domain-domain spacing.

The analyses discussed above provide rich and quantitative metrics of computational actin distributions at length scales similar to experimental actin intensity measurements we have made previously in the TSM1 neuron in the developing Drosophila wing. These metrics allow us to compare values and trends derived for corresponding properties in the computational and experimental distributions and provide strong evidence that the computational model captures fundamental properties of the actin network that exists in a growth cone in vivo. Experimental evidence showed that the growth cone actin network expands progressively over a range of several microns as Abl activity increases, from the Abl-knockdown condition, to wild type, to Abl overexpression (Figure 5D in Clarke *et al;* Clarke *et al*., 2020b), just as the simulations presented here show the multi-micron spread of the actin network with increasing activity of the dominant Abl effector, Arp2/3 (Figure 4). Similarly, quantifying the internal organization of the actin network reveals parallel trends from experiment vs. simulation. The pair-correlation analysis of actin organization performed above queries the internal distribution of actin density in the simulated actin networks in a manner that mirrors the wavelet decomposition of experimental TSM1 growth cone actin distributions we used previously to characterize the internal organization of experimental actin distribution in TSM1. For the in vivo data, the wavelet analysis revealed that the Abl loss-of-function condition induced an enhanced contribution of short length-scale actin spacings to the actin distribution when compared with wild-type (higher-order wavelets; 0.5-4μm spacing. Figure 5 in (Clarke *et al*., 2020b)), consistent with the substantial enhancement of short-range (<2μm) pairwise actin correlations observed computationally at the lowest Arp2/3 levels (Figure 9A, [Arp2/3]<5nM). Conversely, under Abl gain-of-function conditions, wavelet analysis reveals enhanced contribution from long length scales (lower-order wavelets: 4-32μm spacing), consistent with the enhancement of computational pairwise actin correlations >4μm as simulated Arp2/3 levels were increased in the simulations (Please refer to Supplementary Information, Table S3 for more details). Having said this, it is also important to note that it would be an overstatement to call any particular condition from our concentration matrix a unique “model” for the wild type condition of TSM1. Rather, both the simulations here and the quantitative analyses in our previous work (Clarke *et al*., 2020a, 2020b) and (Chandrasekaran et al., submitted) suggest that the wild type condition encompasses substantial stochastic fluctuations that populate a range of actomyosin network architectures. Nonetheless, the quantitative analyses of actin spread and actin internal organization strongly support the conclusion that the trends observed in the computational simulations presented here capture essential properties of the space of actin distributions populated by TSM1 *in vivo* both in wild type and in experimentally perturbed conditions.

Our data suggest that the micron-scale changes to actin networks observed at various levels of Abl signaling can be explained, in part, by filament level changes due to Arp2/3 and Ena. At low Arp2/3 concentrations, Ena-driven polymerase activity alters the filament length distributions slightly towards longer filaments. However, as the Arp2/3 concentration increases, nucleation activity dominates, boosting the abundance of short filaments and reducing both average filament length and the diversity in lengths. As a result, Ena-driven filament length changes saturate in an Arp2/3 dependent manner. The mechanism explaining how the shift in filament distribution to shorter lengths leads to fragmentation of the actin network is presented in detail elsewhere (Chandrasekaran, et al., submitted for publication). In brief, long filaments can bridge between separate domains, allowing them to coalesce under the action of myosin contractility. When short filaments dominate the distribution, the lack of such linking elements causes the network to break down into small, locally contractile domains that are only weakly coupled to one another, as seen here. It is important to note that the concentration bounds of Arp2/3 and Ena used in this work were chosen based on a deterministic model built with published kinetic parameters and were selected before comparing the simulation results with our previous measurements of growth cone actin distributions. Therefore, our observations about the network-level properties of actin reflect inherent emergent properties of the network under standard experimental conditions and were not tuned to match our experimental observations.

While our computational analysis reproduces many properties of Abl function, particularly under WT and overexpressed conditions, the correlation with Abl loss of function is somewhat less complete. At the lowest levels of Arp2/3, simulated actin networks showed condensation of actin, consistent with the actin distribution in Abl loss of function conditions in vivo. However, in the experimental study, we also observe enhanced actin disorder in the most highly condensed actin networks (assayed as Fisher Information Content). This feature was not reproduced in the simulations. The reason for this is not apparent. It may be that a more detailed model of Enabled will illuminate this question. For example, the actin-bundling activity of Ena tetramers has been found crucial in filopodia (Winkelman *et al*., 2014; Harker *et al*., 2019a) and also genetically *in vivo* (Gates *et al*., 2009), so incorporating this activity of Ena in our model may improve the correlation with experimental observations. Additionally, in this study, we investigate the organization of non-polarized actin networks. Polarization aids in directional force generation and may also contribute to the organization of the actin network. Finally, in this study, we have explored the effect of Abl signaling under constant actin conditions, but axon growth is also driven by the dynamic production of actin and other cytoskeletal proteins within the growth cone. Given the evidence that Abl can regulate processes linked to translation (Kannan *et al*., 2014; Kannan and Giniger, 2017), it may be that such effects also modulate Abl-dependent actin organization.

Finally, the critical question motivating this work is how changes in architecture due to Abl signaling may alter functional aspects of the growth cone. While other studies have also employed graph theoretical approaches to characterizing actin organization (Eliaz *et al*., 2020; Floyd *et al*., 2021), the physical and potential biological consequences of such findings have sometimes been elusive. In this study, we designed two perturbative simulations to investigate the effects of Abl-driven changes of actin dynamics in a growth cone-like geometry. The first investigates the transmission of force within static networks built under low-vs. high-Abl-Arp2/3 conditions, while the second investigates the spatial reorganization of fully dynamic low- and high-Abl-Arp2/3 networks in response to perturbation. For the first study, we considered the final axonal architectures from the simulations discussed so far and studied their response to introducing myosin-driven forces exclusively in the densest part of the network. This protocol is akin to published experimental and computational studies that locally activate Myosin 2 in an otherwise frozen actomyosin network(Linsmeier *et al*., 2016). We see that effective transmission of forces across the growth cone occurs under low-Arp2/3 concentrations, while at high-Arp2/3 concentrations, force-transmission is only effective over short distances (sub-micron). This is consistent with the weak structural coupling we observed between actin domains in high-Arp2/3 networks above. For the second study, we restored the complete, force-sensitive mechano-chemical activity of the network to investigate how it responds to short time-scale tensile forces. We embedded mimics of atomic force microscope (AFM) tips functionalized with stable actin filaments in our actin networks and then moved them to exert forces on the network. Here again, we found that at low Arp2/3 levels (equivalent to low Abl), the actin network was displaced as a single unit, while under elevated levels of Arp2/3 (equivalent to enhanced Abl), the weak coupling between actin domains hampers effective force transmission across the network. As a result, only fragments of the overall network move in response to a local pulling force, and the growth cone actin network splits apart. Together, these computational experiments illuminate how the actin cytoskeleton in the growth cone organizes to move as a single entity under wildtype conditions, and perhaps how loss of information- and force-coupling upon Abl overexpression may disrupt growth cone integrity to impair effective guidance.

Our study reveals that Abl signaling can modulate actin organization between connected and fragmented states. We suggest that switching between these states might lend critical mechanistic opportunities to the growth cone as it extends through the embryonic tissue. The connected state helps axons generate and transmit protrusive forces throughout the network, which helps advance the growth cone and keeps it acting as a single unit. On the other hand, one can suppose that growth cone turning requires sampling of information in multiple directions followed by coalescence of structure and force in a single direction. Therefore, we hypothesize that spatial gradients of Abl may create dynamic networks in the growth cone with a range of spatial architectures that switch between globally connected and locally fragmented states to respond directionally to localized extracellular signals.

Axon guidance arises from the action of dozens of neuronal signaling networks, consisting of hundreds or thousands of proteins linked in a wide variety of ways. However, to produce growth cone motility, those signals converge on just a handful of elementary transformations of the actomyosin cytoskeleton. In this work, we have used only the Abl network to manipulate these basic biophysical processes and only one neuron, TSM1, as our biological point of comparison, but the analysis we describe does not depend on those specific choices. Instead, as many cytoskeletal signaling cascades of growth and protrusion terminate on actin polymerization and nucleation as their outputs, we suggest that the mechanistic insights obtained here, especially the fundamental organizational modes, are likely to be broadly relevant to other signaling pathways that regulate growth cone architecture and dynamics. We suggest, therefore, that the perspective we describe here should apply to a wide variety of signaling systems in axon growth and guidance across animal phylogeny.

## Methods

### Mechanochemical Dynamics of Active Matter (MEDYAN)

To computationally simulate actin networks under various levels of Abl, we used MEDYAN(Popov *et al*., 2016), a powerful, freely downloadable (www.medyan.org) active matter simulation software. MEDYAN simulates chemically active reaction networks and enables force-sensitive, stochastic simulations of active filamentous networks, such as actin and microtubules. In MEDYAN, actin filaments are explicitly represented with monomer resolution as a series of non-deformable cylinders coupled at hinge points. Bending deformations around hinge points are allowed corresponding to a persistence length of 16.7µm. (Ott *et al*., 1993) Myosin and α-actinin molecules are represented as Hookean springs with equilibrium lengths based on experimental measurements (Redowicz *et al*., 1999; Ferrer *et al*., 2008). As a result, chemical reactions such as crosslinker (un)binding, motor (un)binding, and motor walking events are defined between filaments that satisfy the appropriate distance thresholds. Branched nucleation is represented as a chemical event leading to the formation of an offspring filament at approximately 70º angle with respect to the parent filament. Ena tetramers are modeled as single molecules that bind to the tip of a free F-actin barbed end and thereby enhancing the polymerase activity.

The simulation space is composed of reaction-diffusion compartments overlaid with the filamentous actin phase. Diffusing reactive molecules (G-actin, unbound crosslinkers, myosins, Arp2/3, and Ena) are considered homogeneously distributed throughout the reaction-diffusion compartments. Thus, a diffusion-reaction is modeled as a hopping reaction between two neighboring compartments. Reaction propensities are defined over the compartment phase, and stochastic trajectories are generated by evolving the chemical reaction network using the *next reaction method*.(Gibson and Bruck, 2000) Chemical events in MEDYAN can lead to mechanical stress accumulation. MEDYAN alternates between short bursts of chemical evolution (25ms) and conjugate gradient mechanical equilibration to generate physically realistic trajectories. Additionally, mechanochemical sensitivity of α-actinin unbinding, myosin walking, and unbinding are incorporated in the model. In this study, we restrict our discussions to actin networks in the growth cone. Please refer to Supplementary Information, Table S1, for details on all parameters used in this study.

## Acknowledgements

We would like to thank all the members of our two labs, particularly Haoran Ni, Qin Ni, Carlos Floyd and Kate O’Neill for their helpful feedback. This work was supported in part by NSF CHE-1800418 and CHE-2102684 and by the Intramural Research Program of NINDS, NIH, grant Z01-NS003013 to EG MEDYAN simulations were carried out on the Deepthought2 Supercomputer at the University of Maryland and Biowulf Supercomputer at the National Institutes of Health.

## Notes

http://www.medyan.org

https://github.com/achansek/readMEDYANtraj

https://github.com/achansek/MEDYANArp23_2021

### Competing Interest Statement

The authors have declared no competing interest.

## References

Araújo, SJ, and Tear, G (2003). Axon guidance mechanisms and molecules: Lessons from invertebrates. Nat Rev Neurosci 4, 910–922.

Bear, JE et al. (2002). Antagonism between Ena/VASP proteins and actin filament capping regulates fibroblast motility. Cell 109, 509–521.

Blondel, VD, Guillaume, JL, Lambiotte, R, and Lefebvre, E (2008). Fast unfolding of communities in large networks. J Stat Mech Theory Exp 2008.

Clarke, A, McQueen, PG, Fang, HY, Kannan, R, Wang, V, McCreedy, E, Buckley, T, Johannessen, E, Wincovitch, S, and Giniger, E (2020a). Dynamic morphogenesis of a pioneer axon in Drosophila and its regulation by Abl tyrosine kinase. Mol Biol Cell 31, 452–465.

Clarke, A, McQueen, PG, Fang, HY, Kannan, R, Wang, V, McCreedy, E, Wincovitch, S, and Giniger, E (2020b). Abl signaling directs growth of a pioneer axon in Drosophila by shaping the intrinsic fluctuations of actin. Mol Biol Cell 31, 466–477.

Dent, EW, and Gertler, FB (2003). Cytoskeletal dynamics and transport in growth cone motility and guidance. Neuron 40, 209–227.

Dent, EW, Gupton, SL, and Gertler, FB (2011). The Growth Cone Cytoskeleton in Axon outgrowth and guidance.pdf. Cold Spring Harb Perspect Biol 3, 1–40.

Eliaz, Y, Nedelec, F, Morrison, G, Levine, H, and Cheung, MS (2020). Insights from graph theory on the morphologies of actomyosin networks with multilinkers. Phys Rev E 102, 1–14.

Feinstein, J, and Ramkhelawon, B (2017). Netrins & Semaphorins: Novel regulators of the immune response. Biochim Biophys Acta -Mol Basis Dis 1863, 3183–3189.

Ferrer, JM, Lee, H, Chen, J, Pelz, B, Nakamura, F, Kamm, RD, and Lang, MJ (2008). Measuring molecular rupture forces between single actin filaments and actin-binding proteins. Proc Natl Acad Sci U S A 105, 9221–9226.

Floyd, C, Levine, H, Jarzynski, C, and Papoian, GA (2021). Understanding cytoskeletal avalanches using mechanical stability analysis. Proc Natl Acad Sci U S A 118, e2110239118.

Floyd, C, Papoian, GA, and Jarzynski, C (2019). Quantifying Dissipation in Actomyosin Networks. Interface Focus 9, 201800781–10.

Forscher, P, and Smith, SJ (1988). Actions of cytochalasins on the organization of actin filaments and microtubules in a neuronal growth cone. J Cell Biol 107, 1505–1516.

Fujiwara, I, Vavylonis, D, and Pollard, TD (2007). Polymerization kinetics of ADP- and ADP-Pi-actin determined by fluorescence microscopy. Proc Natl Acad Sci 104, 8827–8832.

Gallo, G, and Letourneau, PC (2004). Regulation of Growth Cone Actin Filaments by Guidance Cues. J Neurobiol 58, 92–102.

Gates, J, Nowotarski, SH, Yin, H, Mahaffey, JP, Herrera, C, Homem, CCF, Janody, F, Montell, DJ, and Peifer, M (2009). Enabled and Capping protein play important roles in shaping cell behavior during Drosophila oogenesis. Dev Biol 333, 90–107.

Gibson, MA, and Bruck, J (2000). Efficient Exact Stochastic Simulation of Chemical Systems with Many Species and Many Channels. J Phys Chem A 104, 1876–1889.

Giniger, E (2002). How do Rho family GTPases direct axon growth and guidance? A proposal relating signaling pathways to growth cone mechanics. Differentiation 70, 385–396.

van Goor, D, Hyland, C, Schaefer, AW, and Forscher, P (2012). The role of actin turnover in retrograde actin network flow in neuronal growth cones. PLoS One 7.

Grabham, PW, Reznik, B, and Goldberg, DJ (2003). Microtubule and Rac 1-dependent F-actin in growth cones. J. J Cell Sci 116, 3739–3748.

Harker, AJ, Katkar, HH, Bidone, TC, Aydin, F, Voth, GA, Applewhite, DA, and Kovar, DR (2019a). Ena/VASP processive elongation is modulated by avidity on actin filaments bundled by the filopodia cross-linker fascin. Mol Biol Cell 30, 851–862.

Harker, AJ, Katkar, HH, Bidone, TC, Aydin, F, Voth, GA, Applewhite, DA, and Kovar, DR (2019b). Ena/VASP processive elongation is modulated by avidity on actin filaments bundled by the filopodia crosslinker fascin. Mol Biol Cell 30, mbc.E18-08-0500.

Hinck, L (2004). The versatile roles of “axon guidance” cues in tissue morphogenesis. Dev Cell 7, 783–793.

Huber, AB, Kolodkin, AL, Ginty, DD, and Cloutier, JF (2003). Signaling at the growth cone: Ligand-receptor complexes and the control of axon growth and guidance. Annu Rev Neurosci 26, 509–563.

Jones, SB, Lu, HY, and Lu, Q (2004). Abl tyrosine kinase promotes dendrogenesis by inducing actin cytoskeletal rearrangements in cooperation with Rho family small GTPases in hippocampal neurons. J Neurosci 24, 8510–8521.

Kahn, OI, and Baas, PW (2016). Microtubules and Growth Cones: Motors Drive the Turn. Trends Neurosci 39, 433–440.

Kannan, R et al. (2017). The Abl pathway bifurcates to balance Enabled and Rac signaling in axon patterning in Drosophila. Development 144, 487–498.

Kannan, R, and Giniger, E (2017). New perspectives on the roles of Abl tyrosine kinase in axon patterning. Fly (Austin) 0, 1–11.

Kannan, R, Kuzina, I, Wincovitch, S, and Nowotarski, SH (2014). The Abl / Enabled signaling pathway regulates Golgi architecture in Drosophila photoreceptor neurons. Mol Biol Cell 25, 2993–3002.

Li, X, Ni, Q, He, X, Kong, J, Lim, S-M, Papoian, GA, Trzeciakowski, JP, Trache, A, and Jiang, Y (2020). Tensile force-induced cytoskeletal remodeling: Mechanics before chemistry. PLOS Comput Biol 16, e1007693.

Linsmeier, I, Banerjee, S, Oakes, PW, Jung, W, Kim, T, and Murrell, MP (2016). Disordered actomyosin networks are sufficient to produce cooperative and telescopic contractility. Nat Commun 7, 12615.

Lowery, LA, and Vactor, D Van (2009). The trip of the tip: Understanding the growth cone machinery. Nat Rev Mol Cell Biol 10, 332–343.

Mogilner, a, and Oster, G (1996). Cell motility driven by actin polymerization. Biophys J 71, 3030–3045.

Mullins, RD, Heuser, JA, and Pollard, TD (1998). The interaction of Arp2/3 complex with actin: Nucleation, high affinity pointed end capping, and formation of branching networks of filaments. Proc Natl Acad Sci 95, 6181–6186.

Ni, Q, and Papoian, GA (2019). Turnover versus Treadmilling in Actin Network Assembly and Remodeling. Cytoskeleton 76, 562–570.

O’Connor, TP, Duerr, JS, and Bentley, D (1990). Pioneer growth cone steering decisions mediated by single filopodial contacts in situ. J Neurosci 10, 3935–3946.

Omotade, OF, Pollitt, SL, and Zheng, JQ (2017). Actin-based growth cone motility and guidance. Mol Cell Neurosci 84, 4–10.

Ott, A, Magnasco, M, Simon, A, and Libchaber, A (1993). Measurement of the persistence length of polymerized actin using fluorescence microscopy. Phys Rev E 48.

Pandit, NG, Cao, W, Bibeau, J, Johnson-chavarria, EM, and Taylor, EW (2020). Force and phosphate release from Arp2 / 3 complex promote dissociation of actin filament branches. Proc Natl Acad Sci.

Pollard, TD (2007). Regulation of actin filament assembly by Arp2/3 complex and formins. Annu Rev Biophys Biomol Struct 36, 451–477.

Popov, K, Komianos, J, and Papoian, GA (2016). MEDYAN : Mechanochemical Simulations of Contraction and Polarity Alignment in Actomyosin Networks. PLoS Comput Biol 12, e1004877.

Redowicz, MJ, Hammer, JA, Bowers, B, Zolkiewski, M, Ginsburg, A, Korn, ED, and Rau, DC (1999). Flexibility of Acanthamoeba myosin rod minifilaments. Biochemistry 38, 7243–7252.

Sánchez-Soriano, N, Gonçalves-Pimentel, C, Beaven, R, Haessler, U, Ofner-Ziegenfuss, L, Ballestrem, C, and Prokop, A (2010). Drosophila growth cones: A genetically tractable platform for the analysis of axonal growth dynamics. Dev Neurobiol 70, 58–71.

Santos, TE, Schaffran, B, Broguière, N, Meyn, L, Zenobi-Wong, M, and Bradke, F (2020). Axon Growth of CNS Neurons in Three Dimensions Is Amoeboid and Independent of Adhesions. Cell Rep 32.

Schaefer, AW, Kabir, N, and Forscher, P (2002). Filopodia and actin arcs guide the assembly and transport of two populations of microtubules with unique dynamic parameters in neuronal growth cones. J Cell Biol 158, 139–152.

Seiradake, E, Jones, EY, and Klein, R (2016). Structural Perspectives on Axon Guidance.

Strasser, GA, Rahim, NA, Vanderwaal, KE, Gertler, FB, and Lanier, LM (2004). Arp2/3 is a negative regulator of growth cone translocation. Neuron 43, 81–94.

Suter, DM, and Forscher, P (2010). Substrate–Cytoskeletal Coupling as a Mechanism for the Regulation of Growth Cone Motility and Guidance. J Neurobiol 44, 97–113.

Tamariz, E, and Varela-Echavarría, A (2015). The discovery of the growth cone and its influence on the study of axon guidance. Front Neuroanat 9, 1–9.

Turney, SG, Ahmed, M, Chandrasekar, I, Wysolmerski, RB, Goeckeler, ZM, Rioux, RM, Whitesides, GM, and Bridgman, PC (2016). Nerve growth factor stimulates axon outgrowth through negative regulation of growth cone actomyosin restraint of microtubule advance. Mol Biol Cell 27, 500–517.

Winkelman, JD, Bilancia, CG, Peifer, M, and Kovar, DR (2014). Ena/VASP Enabled is a highly processive actin polymerase tailored to self-assemble parallel-bundled F-actin networks with Fascin. Proc Natl Acad Sci 111, 4121–4126.

Witte, H, and Bradke, F (2008). The role of the cytoskeleton during neuronal polarization. Curr Opin Neurobiol 18, 479–487.

