## Supplemental tables and figures for "Computational simulations reveal that Abl activity controls cohesiveness of actin networks in growth cones"

### Supplementary Information

| Parameter | Symbol | Value | Reference |
|---|---|---|---|
| **Geometric parameters** | | | |
| Compartment size | $L_{comp}$ | 500nm | [1] |
| Number of compartments in each dimension | Nx,Ny,Nz | 4,4,15 | - |
| Length of a cylinder | $L_{cyl}$ | 40 | - |
| Minifilament binding distance | $d_{NMII,bind}$ | 175-225nm | [1] |
| α-actinin binding distance | $d_{\alpha,bind}$ | 30-40nm | [1] |
| **Diffusion rates** | | | |
| Actin | $k_{actin,diff}$ | 20μm²/s [80 $s^{-1}$] | [1] |
| α-actinin | $k_{\alpha,diff}$ | $k_{actin,diff}/10$ | [1] |
| Myosin minifilament | $k_{NMII,diff}$ | $k_{actin,diff}/100$ | [1] |
| Enabled | $k_{Ena,diff}$ | $k_{actin,diff}/100$ | - |
| Arp2/3 | $k_{Arp,diff}$ | $k_{actin,diff}/100$ | - |
| **Kinetic rate constants** | | | |
| Actin polymerization at plus end | $k_{actin\ poly,+}$ | 11.6 $(\mu M.s)^{-1}$ [0.154$s^{-1}$] | [2] |
| Actin **de**polymerization at plus end | $k_{actin\ depoly,+}$ | 1.3 $(\mu M.s)^{-1}$ [0.017$s^{-1}$] | [2] |
| Actin polymerization at minus-end | $k_{actin\ poly,-}$ | 1.4 $s^{-1}$ | [2] |
| Actin **de**polymerization at minus end | $k_{actin\ depoly,-}$ | 0.8 $s^{-1}$ | [2] |
| α-actinin binding | $k_{\alpha,bind}$ | 0.7 $(\mu M.s)^{-1}$ [0.009 $s^{-1}$] | [3] |
| α-actinin unbinding (F=0pN) | $k_{\alpha,unbind}$ | 0.3 $s^{-1}$ | [3] |
| NMII head binding | $k_{NMII,bind}$ | 0.2$s^{-1}$ | [4] |
| NMII head unbinding (F=0pN) | $k_{NMII,unbind}$ | 1.0$s^{-1}$ | **a** |
| Arp2/3 binding | $k_{Arp,bind}$ | 0.0017$s^{-1}$ | **b**[5] |
| Arp2/3 unbinding (F=0pN) | $k_{Arp,unbind}$ | 0.02$s^{-1}$ | **c**[5] |
| Ena binding | $k_{Ena,bind}$ | 75 $(\mu M.s)^{-1}$ [0.996$s^{-1}$] | [6] |
| Ena unbinding | $k_{Ena,unbind}$ | 0.69$s^{-1}$ | [6] |
| Ena enhanced polymerization | $k_{Ena,poly}$ | 29.38 $(\mu M.s)^{-1}$ [0.39$s^{-1}$] | **d** |
| **Mechanochemical constants** | | | |
| NMII/α-actinin binding pitch on actin filament | - | 27nm | [1] |
| NMII head step size | $d_{step}$ | 6.0nm | **e**[7][8] |
| NMII minifilament range of number of heads on each side of bipolar minifilament | $N_{min}$-$N_{max}$ | 15-30 | **f**[9,10] |
| NMII minifilament stall force | $F_{NMII,stall}$ | 300pN for minifilament | **g** |

| | | | |
|---|---|---|---|
| NMII per head unbinding Force | $F_{NMII,\ unbind}$ | 12.62pN per head | [11] |
| Tunable parameters | $\beta$ | 0.2 | [1] |
| | $\gamma$ | $0.05pN^{-1}$ | [1] |
| | $\zeta$ | 0.1 | [1] |
| Linker unbinding force | $F_{\alpha,\ unbind}$ | 17.2pN | [12] |
| Characteristic force of Brownian ratchet | $F_{ratchet}$ | 1.5pN | [13] |
| **Mechanical constants** | | | |
| Actin filament stretching constant | $K_{fil,str}$ | 100pN/nm | [1] |
| Actin filament bending energy | $\varepsilon_{bend}$ | 2690pN.nm | [1] |
| Cylinder-Cylinder Excluded volume constant | $K_{vol}$ | $10^5$pN/nm | [1] |
| Myosin cross-bridge stiffness | $K_{NMII,str}$ | 2.5pN/nm | [8] |
| α-actinin stiffness | $K_{\alpha,str}$ | 8pN/nm | [14] |
| Boundary repulsion energy | $\varepsilon_{boundary}$ | $10k_BT$ | - |
| Boundary repulsion screening length | $\Lambda$ | 2.7nm | - |
| Minimization parameters | | | |
| Length of the chemical step | $\delta_{chemistry}$ | 25ms | - |
| Force tolerance for mechanical minimization | $F_T$ | 10pN | - |

**Supplementary Table S1 Table of simulation parameters used in MEDYANv4.1 to simulate dendritic actin networks.**

**a**Obtained by assuming a duty ratio of 17%, which is the average of the 11% duty ratio corresponding to NMIIA and 23% duty ratio corresponding to NMIIB.[4]

**b**Slope of Figure 2D.

**c**Based on ATP actin parent filaments.

**d** Refer Supporting Information, Section A.1

**e**Value chosen is close to the *Dictyostelium* step size of 7.3±0.4nm

**f** Experimental results suggest total number of heads in NMIIA- 56[10], 58[9], NMIIB-60[9], and NMIIC-28[9]. Here, we use a wide range to account for multiple binding modes of myosin isoforms.

**g**Considering the myosin-actin cross-bridge stiffness of $k_{mhead} = 2.5$pN/nm [8], the stall force of a single head $F_{head} = k_{mhead} x d_{step}$=15pN. If 15 heads are bound, the $F_{stall,15}$=225pN, while 30 bound heads result in $F_{stall,30}$=450pN. As our simulations dynamically choose myosin heads at binding, a stall force of about 300pN was chosen.

| Perturbation study paramters | Symbol | Value | Reference |
|---|---|---|---|
| **Myosin walking driven perturbation of network** | | | |
| Length of the chemical step | $\delta_{chemistry}$ | 10ms | - |
| Snapshot time step | $\Delta_{snapshot}$ | 100ms | - |
| Width of the active zone where myosin is activated | $d_{activezone}$ | 500nm | - |
| **Functionalized AFM probe mimic simulations** | | | |
| Number of compartments in each dimension | Nx,Ny,Nz | 4,4,19 | - |
| The time scale of the chemical step | $\delta_{chemistry}$ | 25ms | - |
| AFM tip motion at the end of the chemical cycle | $d_{AFM}$ | 5nm | [15] |
| Total number of AFM tip displacements | $N_{AFMsteps}$ | 100 | - |
| Effective AFM tip velocity | $v_{AFM}$ | 200nm/s | - |
| AFM tip radius | $R_{AFM}$ | 250nm | [15] |
| Number of filaments attached to AFM tip | $N_{filaments,AFM}$ | 50 | [15] |
| Length of filaments attached to AFM tip | $L_{filaments,AFM}$ | 540nm | - |

**Supplementary Table S2 Table of additional simulation parameters used in MEDYANv4.1 to study responses of actin networks to perturbations.**

| Property/measurable | Experimental observation | Computational result |
|---|---|---|
| **Actin spread (σ)** | Actin profile spread - $\sigma_{Abl\ LOF} < \sigma_{WTAbl} < \sigma_{Abl\ GOF}$ | Simulation profile spread - Spread increases along Arp axis over larger length scales than Ena axis |
| **Actin organization** | Wavelet analysis – measures spatial actin organization pattern AblLOF profiles have increased contribution from low order wavelets (~0.5-4μm) * AblGOF profiles have increased contribution from higher order wavelets (~4-32μm) * | Pair correlation function – As Arp2/3 is increased, g(r)=1 (bulk density = local density) at longer length scales. |
| **Actin disorder/fragmentation** | Fisher information reveals spatial actin disorder under both Abl LOF and Abl GOF | Louvain modularity optimization - As Arp2/3 is increased, the number of communities increases Two perturbative simulations reveal actin network is mechano-chemically fragmented at [Arp2/3]≥10nM |

**Supplementary Table S3 Table showing results from live imaging of Abl axons under genetic mutation against results from simulations in this study.** Experimental results mentioned are from Clarke et al. [16,17].

* For comparing pair-correlation results with the wavelet analysis of previous experimental data (Figure 5 in Clarke et al, 2020 [17]), note that the calculation of a given wavelet order includes the terms for the next two higher orders. Thus, calculation of the order 7 wavelet (corresponding to data with a nominal spacing of 2-4μm in the experimental dataset) incorporates the contributions from orders 8 and 9 (nominal spacing 1-2 and 0.5-1μm, respectively), while order 4 (16-32μm) incorporates the terms corresponding to 8-16 and 4-8μm.

**Supplementary Figures**

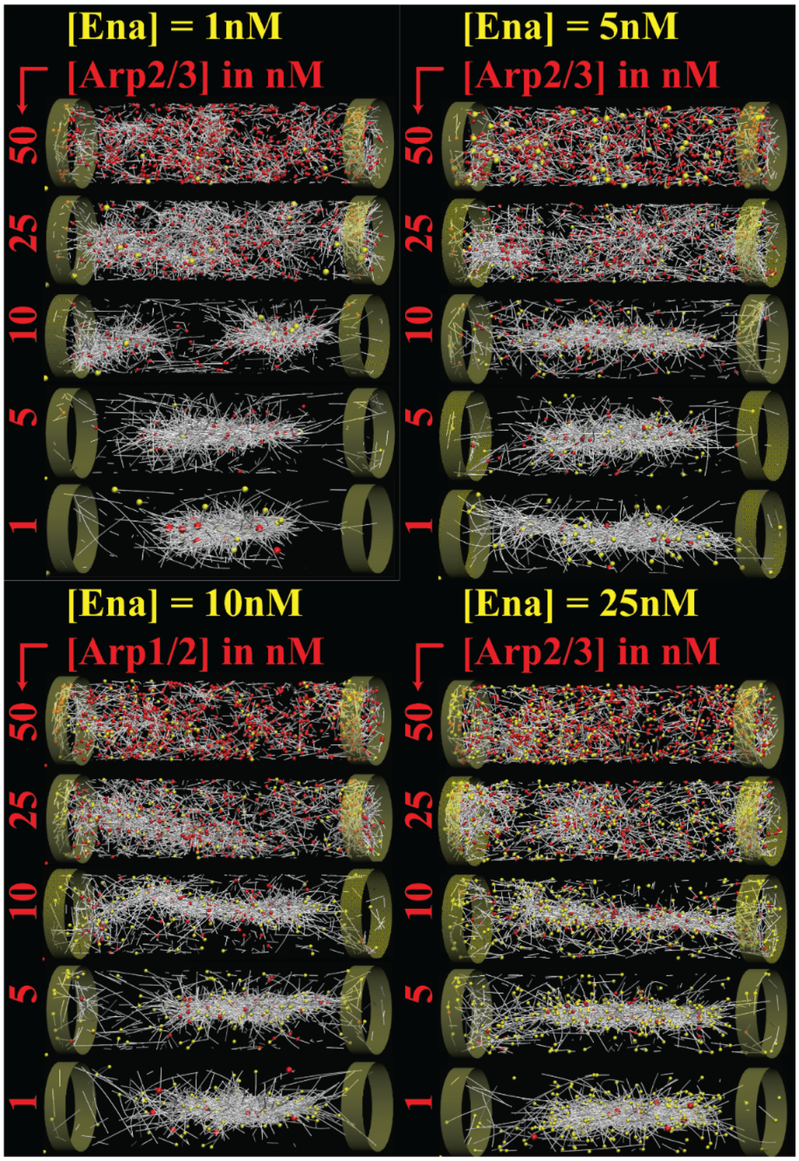


**Supplementary Figure S1 Representative final snapshots showing actin networks at various [Arp2/3] concentrations at [Ena]=1nM, 5nM, 10nM and 25nM.** Actin filaments, Arp2/3, and Ena are shown as white filaments, red spheres, and yellow spheres, respectively.

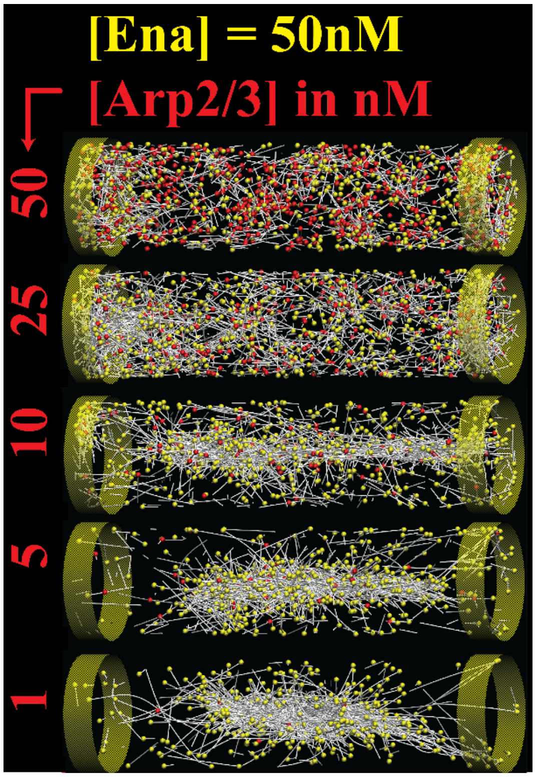


**Supplementary Figure S2 Representative final snapshots showing actin networks at various [Arp2/3] concentrations at [Ena]=50nM.** Actin filaments, Arp2/3, and Ena are shown as white filaments, red spheres, and yellow spheres, respectively.

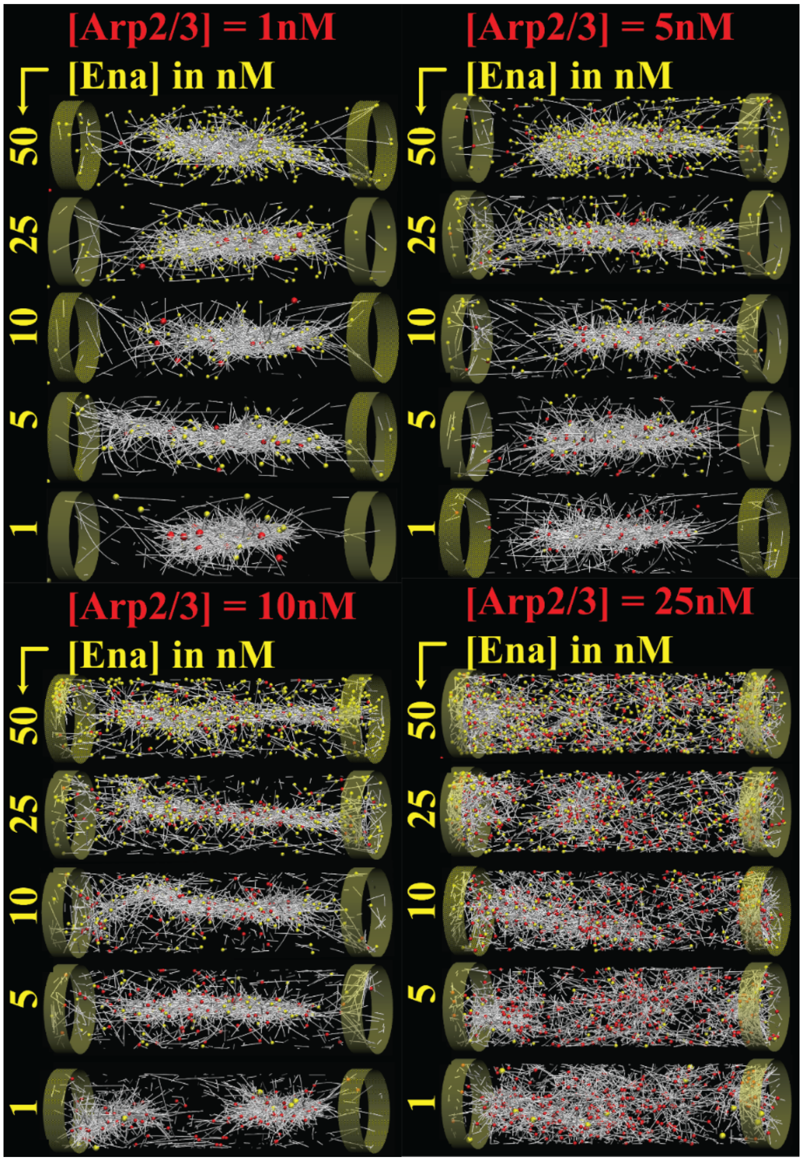


**Supplementary Figure S3 Representative final snapshots showing actin networks at various [Ena] concentrations at [Arp2/3]=1nM, 5nM, 10nM and 25nM.** Actin filaments, Arp2/3, and Ena are shown as white filaments, red spheres, and yellow spheres, respectively.

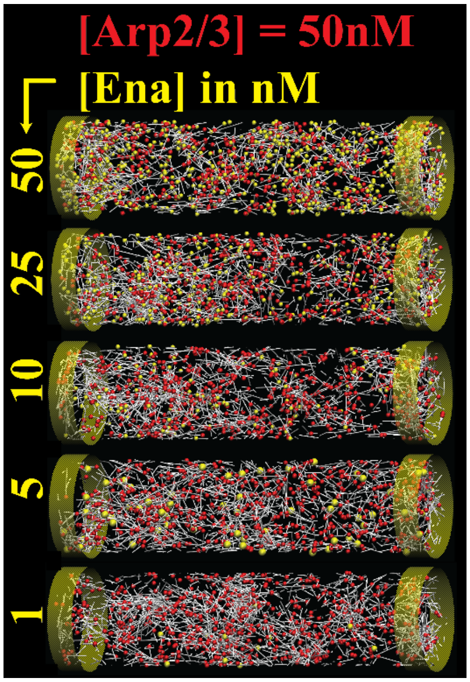


**Supplementary Figure S4 Representative final snapshots showing actin networks at various [Ena] concentrations at [Arp2/3]=50nM.** Actin filaments, Arp2/3, and Ena are shown as white filaments, red spheres, and yellow spheres, respectively.

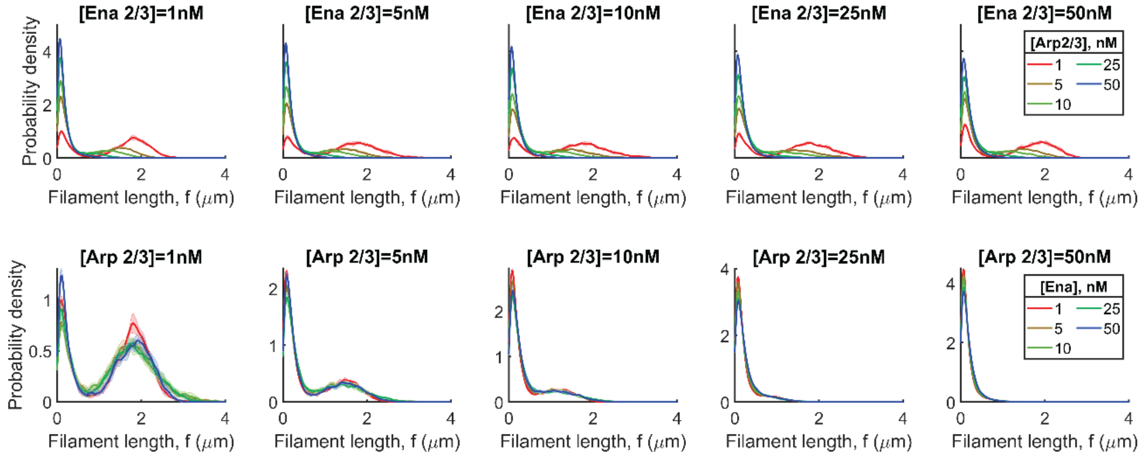


**Supplementary Figure S5 Effect of Arp2/3 and Ena on filament length distributions.** Probability density function profiles of filament length distributions are shown by varying [Ena] concentration along the top row and by varying [Arp2/3] concentration along the bottom row. Each panel in the top row shows profiles from different Arp2/3 concentrations (shown in legend), while each panel in the bottom row shows profiles from varying Ena concentrations. Solid lines and shaded areas represent mean and standard deviation, respectively.

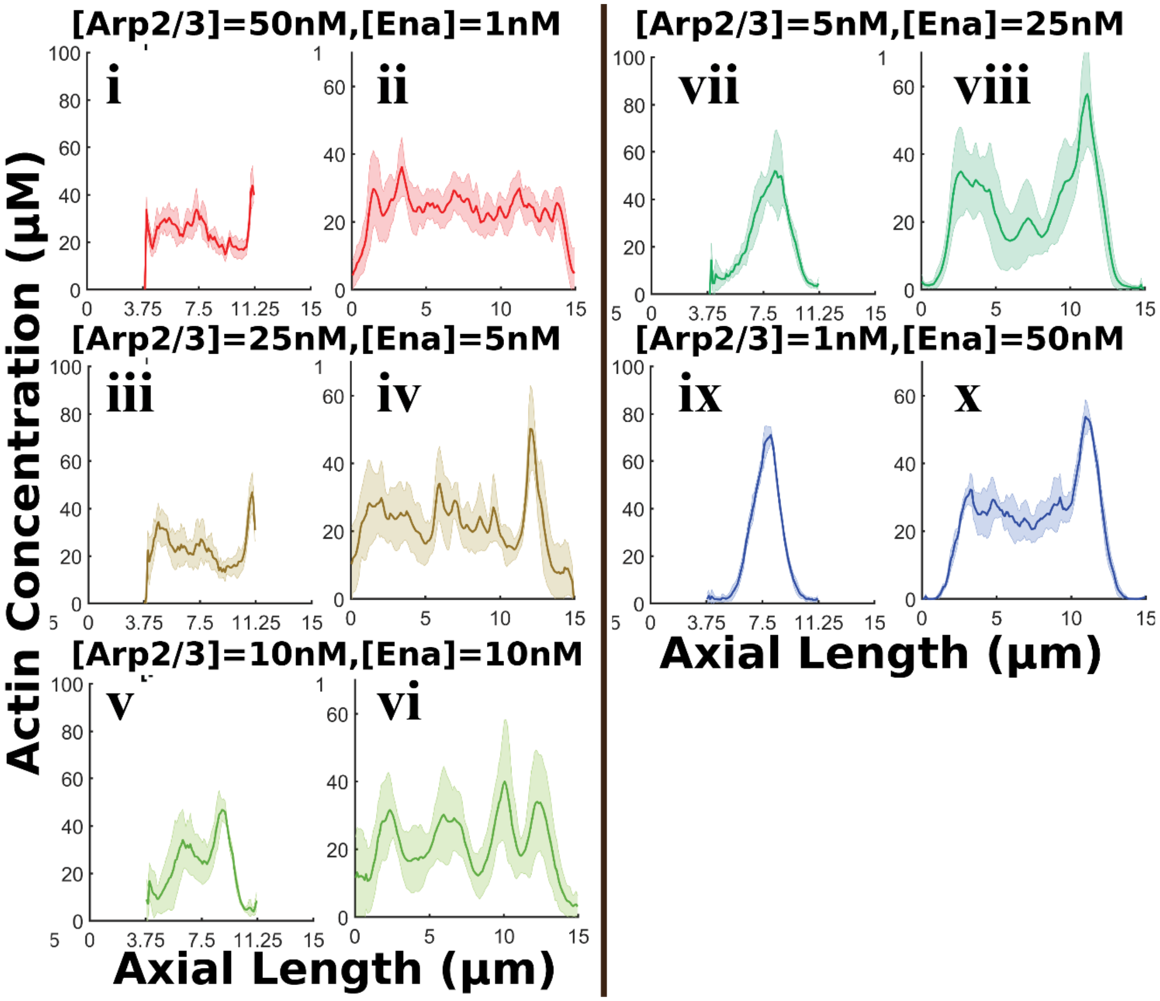


**Supplementary Figure S6 Simulations at longer length scales preserve salient features found at lower length scales.** In each row, the left panel (i,iii,v,vii,x) shows peak-aligned actin profiles from simulations in 7.5μm long reaction volumes, while the right panel (ii, iv, vi, viii, x) shows peak-aligned actin profiles from simulations in 15μm long reaction volumes. The peak-aligned profiles from 7.5 μm long reaction volumes were translated appropriately to enable easy comparison to profiles obtained from simulations in 15μm reaction volume.

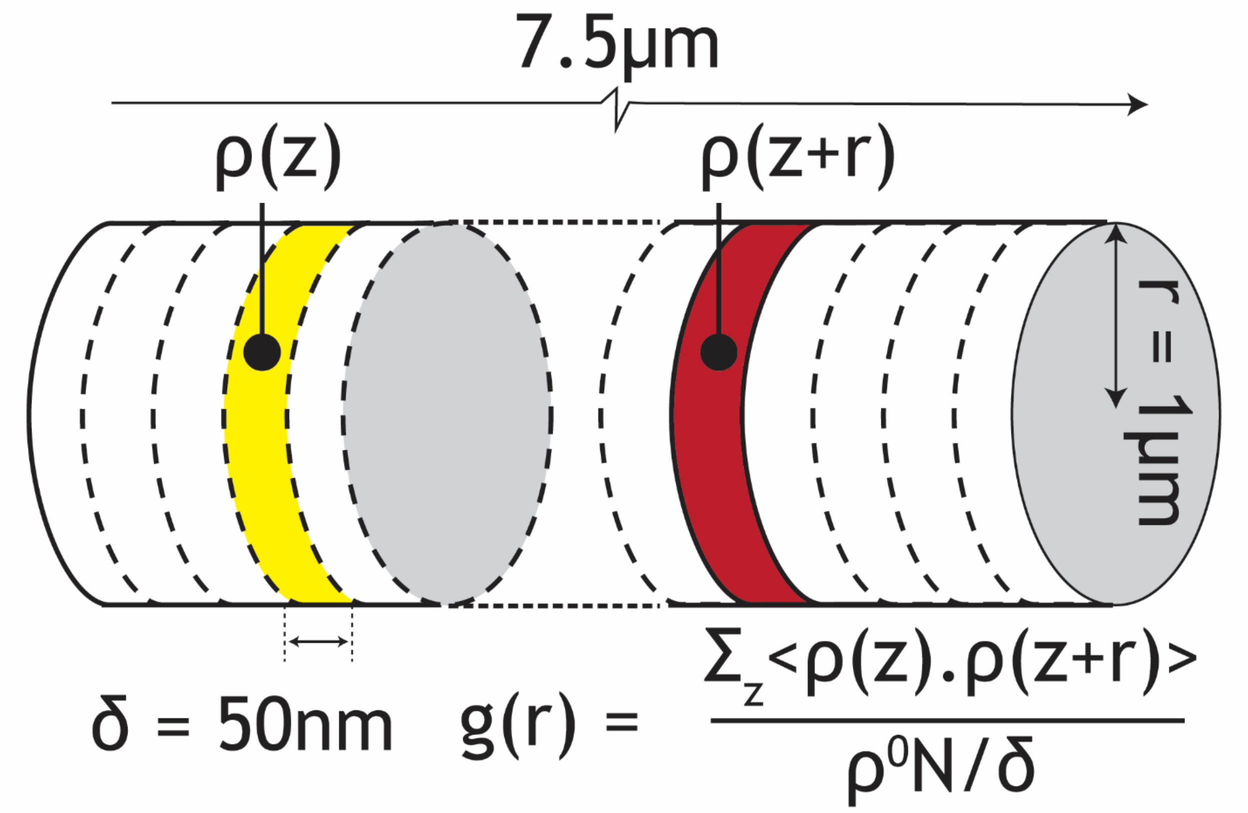


**Supplementary Figure S7 Schematic for calculating pair correlation function, g(r).** Reaction volume consists of N F-actin monomers and is divided into sub-cylinders of width δ=50 nm. The linear density, defined as the number of F-actin monomers per unit length is defined for each sub-cylinder (ρ(z)) and the reaction volume ($\rho^0$). Angular brackets represent averaging over multiple snapshots and replicates.

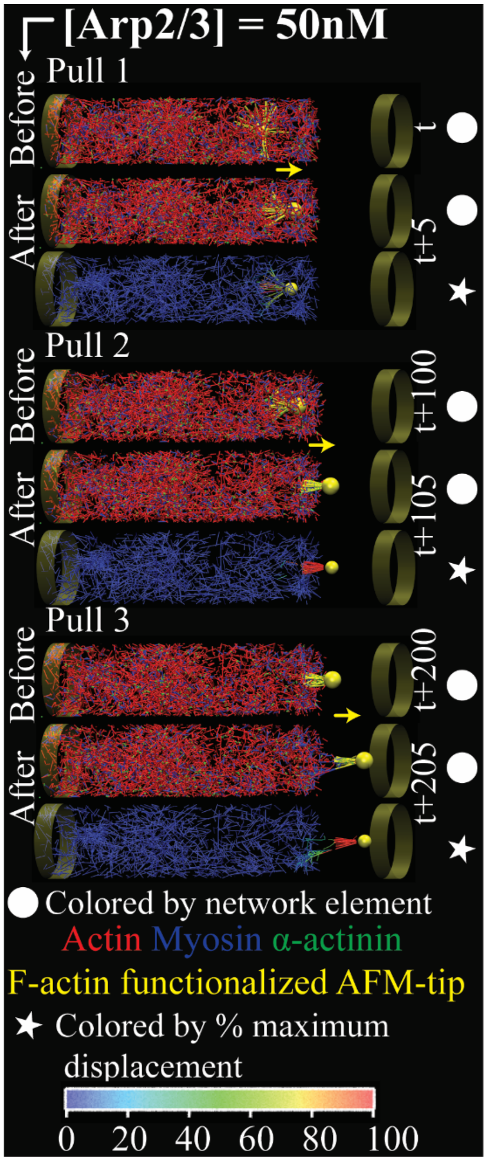


**Supplementary Figure S8 Mechanical perturbation by AFM probe reveals mechanochemical fragmentation of [Arp2/3]=50nM networks.** Three snapshots are shown corresponding to each AFM probe pull event. For each event, the snapshots before and after AFM tip displacement are visualized showing actin (red), myosin (blue), α-actinin (green) and the AFM probe tip (yellow sphere) and the actin filaments attached to it (yellow filaments). Yellow arrows show the direction of tip displacement. Finally, we also visualize the filaments after AFM probe displacement and color each filament based on their displacement during the pull event (color bar shown below the image).

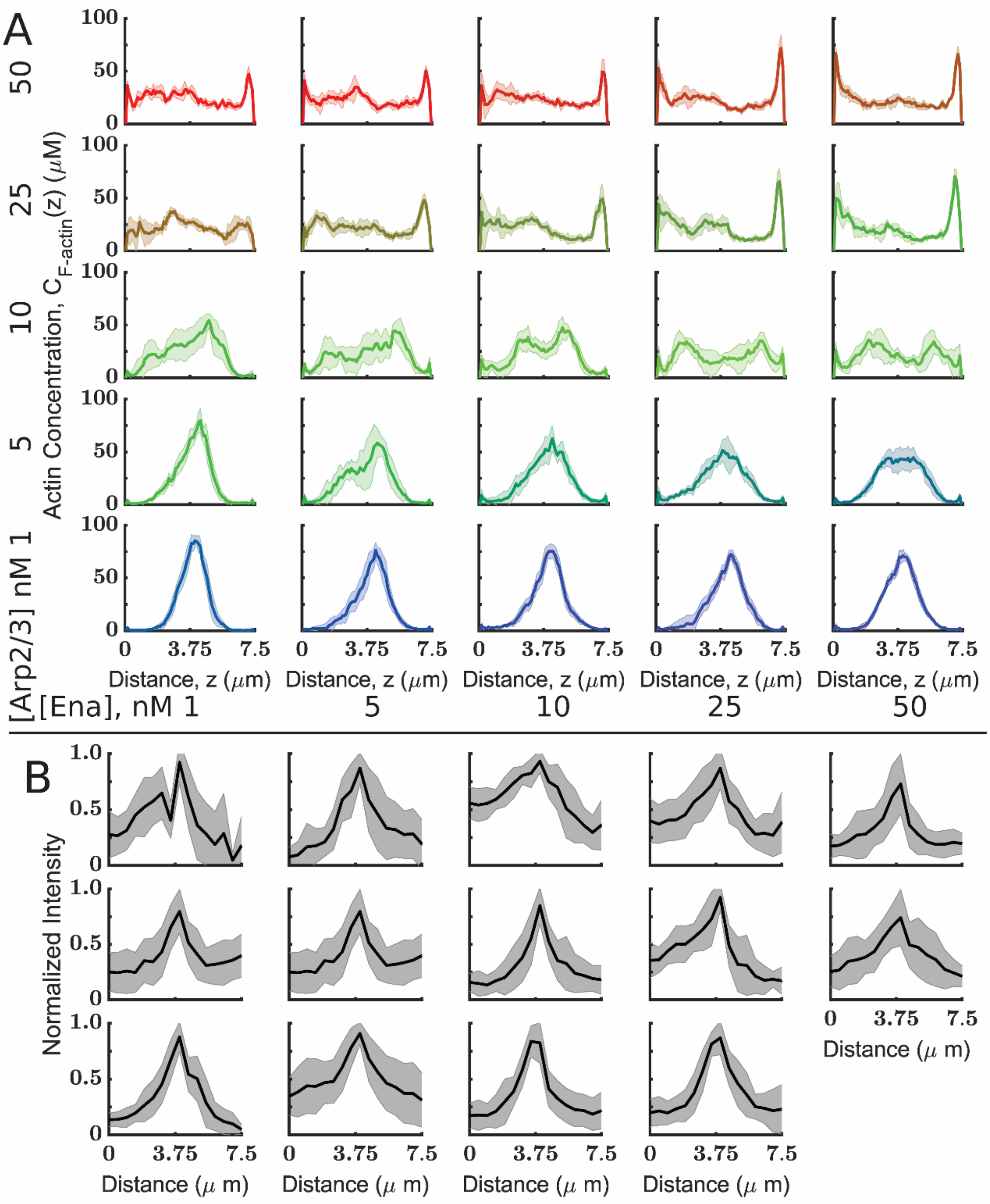


**Supplementary Figure S9 Simulations capture features of axonal growth cones observed in experiments.** A. Peak-aligned actin concentrations are plotted as a function of distance along the cylinder axis. Arp2/3 concentrations are mentioned to the left, while Ena concentrations are labeled at the bottom of the panel. The solid line and shaded area represent mean and standard deviation, respectively. Note that these are the same curves displayed in the lower portion of 3, reproduced here to facilitate comparison to Panel B. B. Actin intensity distribution in 14 WT Abl axons. Each sub-panel shows the mean and standard deviation of peak-aligned actin profiles from the first 30 minutes of imaging a WT axon. Experimental actin intensity distributions were truncated 3.75 µm on either side of the peak according to experimental determination of growth cone width. Please note that the peak of experimental distributions was set to 3.75µm to aid easy comparison with the simulation data.

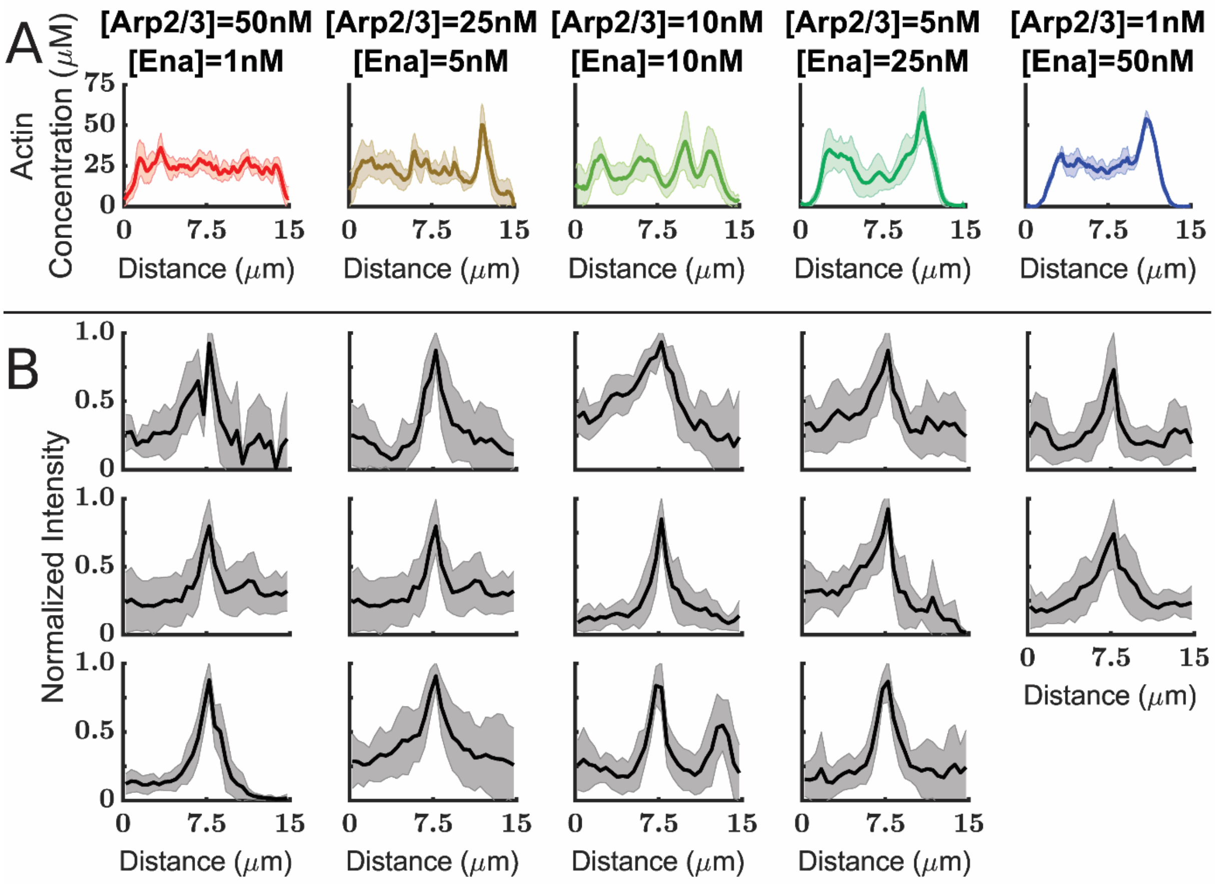


**Supplementary Figure S10 In-silico mimics of Abl in 15µm simulations share similar properties with *in vivo* growth cones.** A. Axial actin distribution from the final snapshot of simulations is shown at various Arp/3 and Ena concentrations. Each panel shows actin peak-aligned actin profiles from 6 replicates. B. Peak-aligned, normalized actin intensity profiles from WT Abl axons is shown. The actin profiles are truncated based on an experimentally determined average growth cone span of 15µm. Each panel shows actin profiles from the first 30 minutes of imaging (sampling frequency=3min) as solid lines. In all 14 cells imaged, the mean and standard deviation of the actin intensity profiles are shown as solid black lines and shaded areas.

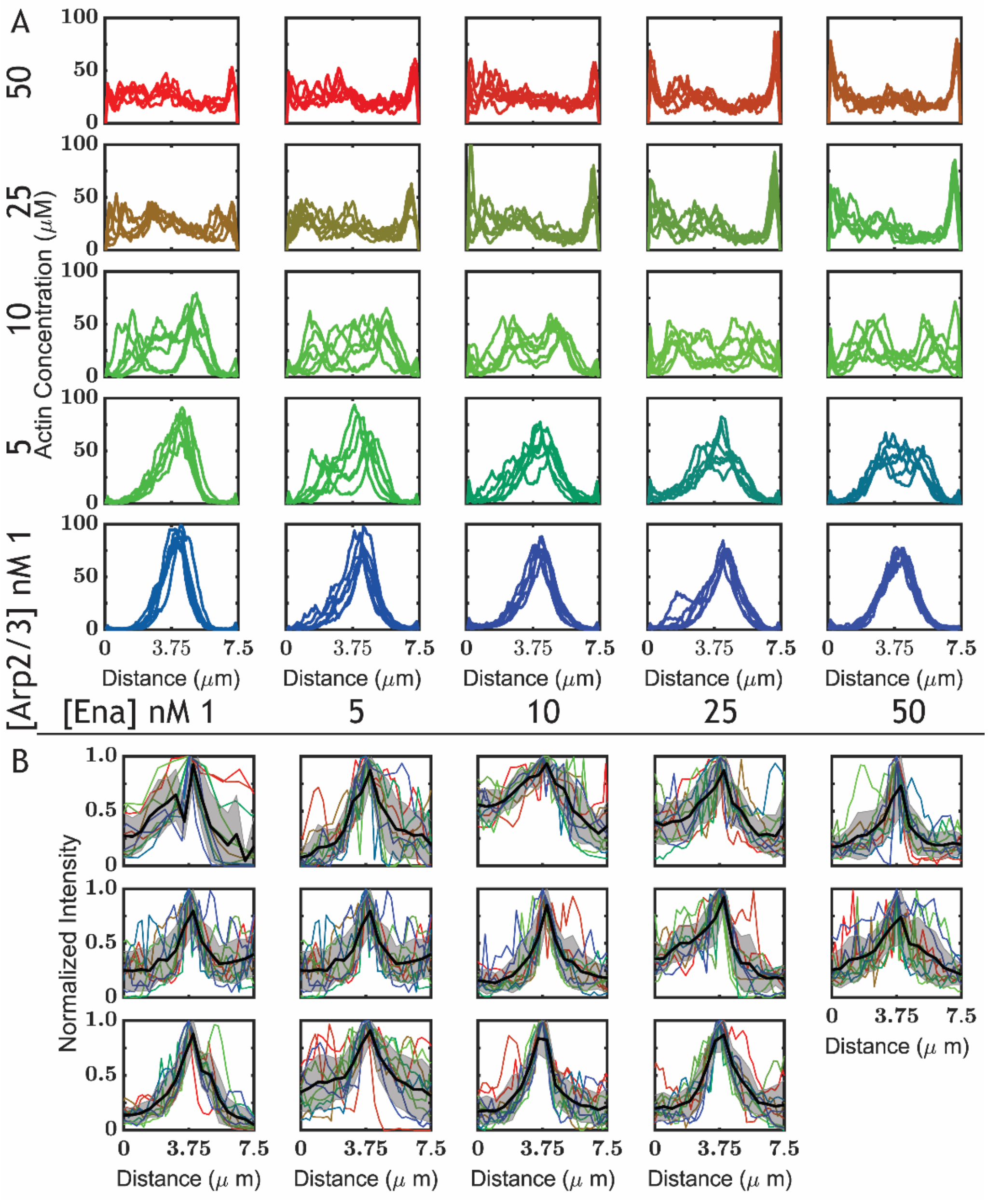


**Supplementary Figure S11 In-silico mimics of Abl share similar properties with *in vivo* growth cones.** A. Axial actin distribution from the final snapshot of simulations is shown at various Arp/3 and Ena concentrations. Each panel shows actin peak-aligned actin profiles from 6 replicates. B. Peak-aligned, normalized actin intensity profiles from wild type axons is shown. The actin profiles are truncated at a span of 7.5μm. Each panel shows actin profiles from the first 30 minutes of imaging (sampling frequency=3min) as solid lines. In all 14 cells imaged, the mean and standard deviation of the actin intensity profiles are shown as solid black lines and shaded areas.

**Supplementary Methods**

A.1. Determination of Enabled-enhanced polymerization rate

In our simulations, we model Enabled as a molecule that can bind free barbed ends to stabilize them from depolymerization. Additionally, Ena also enhances the polymerization rate of filaments. As Ena polymerization rates are not readily available, we used data from Winkelman et al.[18] to determine rate constants. Under 0.924μM actin, 5nM of Ena extends actin filaments at the rate of 27.1 subunits/(filament.s) in the first 150s. The filament extension rate ($\frac{dN_{EV}^{+}}{dt}$) of Ena bound filaments, can be written as,

$$\frac{dN_{EV}^{+}}{dt} = k_{EV,poly}^{+} C_{A}^{o}$$

Thus, the Ena driven elongation rate of barbed ends is given as, $k_{EV,poly}^{+} = 29.38 \mu M^{-1} s^{-1}$.

A.2. A deterministic model for actin filament dynamics in the presence of Arp2/3 and Ena

A deterministic ODE model of Arp2/3 and Ena dynamics was used to determine bounds on Arp2/3 and Ena concentrations. Please refer to Appendix Table C-1 for a list of notations used. The following table gives the concentration of species

| **Concentration of** | **Symbol** |
|---|---|
| F-actin | [FA] |
| Minus End | [M] |
| Plus End | [P] |
| Filament bound Arp2/3 | [B] |
| Ena bound plus end | [EP] |
| Diffusing Actin | [AD] |
| Diffusing Arp2/3 | [BD] |
| Diffusing Ena | [ED] |

**Table S4 Table of concentrations used in the deterministic model of actin dynamics.**

The following kinetic equations give the dynamics of actin under the presence of Arp2/3 and Ena.

$$\frac{d[FA]}{dt} = k_{Arp,bind}\frac{1}{n}[FA][AD][BD] + k_{poly,+}[AD][P] + k_{poly,-}[AD][M] - k_{depoly,-}[AD][M]$$

$$- k_{depoly,+}[P] - k_{depoly,-}[M] + k_{poly,Ena}[EP][AD]$$

$$\frac{d[AD]}{dt} = -\frac{d[FA]}{dt}$$

$$\frac{d[EP]}{dt} = k_{bind,Ena}[P][ED] - k_{unbind,Ena}[EP]$$

$$\frac{d[P]}{dt} = -\frac{d[EP]}{dt} + k_{Arp,bind}\frac{1}{n}[FA][AD][BD]$$

$$\frac{d[M]}{dt} = k_{Arp,unbind}[B]$$

$$\frac{d[B]}{dt} = k_{Arp,bind}\frac{1}{n}[FA][AD][BD] - k_{Arp,unbind}[B]$$

$$\frac{d[BD]}{dt} = -\frac{d[B]}{dt}$$

The equations were solved simultaneously in MATLAB® using the ode45 function to generate numerical profiles of average filament length and Ena-driven actin turnover. Figure S12 shows limited changes to filament lengths by Arp2/3 and the total amount of actin polymerized by Ena above 50nM concentrations. Hence, we simulated networks at concentrations of 1, 5, 10, 25, and 50nM Arp2/3 and Ena.

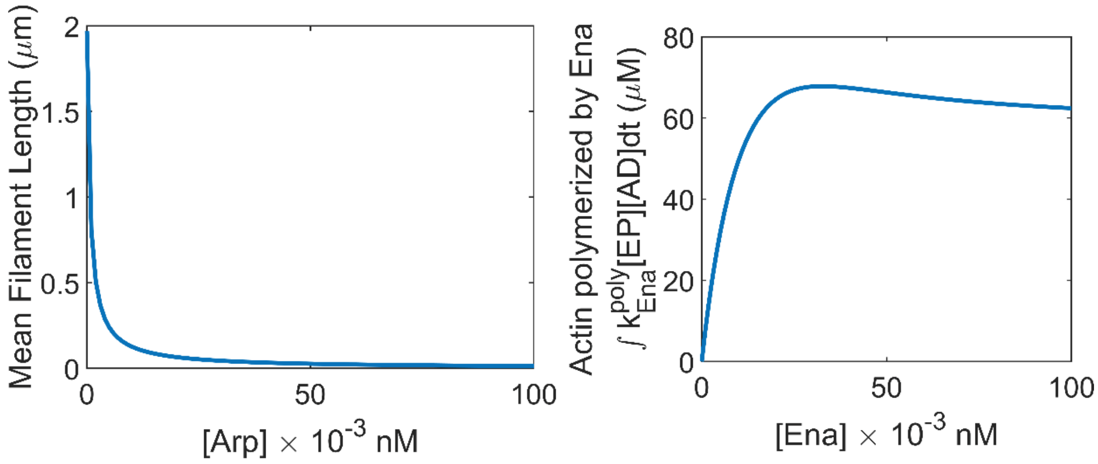

**Supplementary Figure S12 Deterministic model for branching and Ena kinetics predicts bounds for Arp2/3 and Ena concentrations.** A. Plot of mean filament length predicted by a deterministic model of filament dynamics at various Arp2/3 concentrations ([Ena]=0nM]). B. Plot of total diffusing actin that is converted to filamentous actin through Ena-driven filament extension at various Ena concentrations ([Arp2/3] = 0nM).

A.3. Filament-Filament contacts based community detection using Louvain Modularity

Modularity metric, Q$\epsilon$[-1,1], measures how well a set of edge-weighted nodes are organized as communities.

$$Q = \frac{1}{2m}\sum_{i,j}\left[A_{ij} - \frac{k_i k_j}{2m}\right]\delta(c_i, c_j)$$

$A_{i,j}$ represents the edge weight between nodes i and j. $k_i$ and $k_j$ represent the sum of weights from all nodes that are connected to i and j, respectively. ci and cj represent communities that nodes i and j are assigned to. Finally, $m = \frac{1}{2}\sum A_{ij}$. Louvain modularity optimization algorithm was implemented in MATLAB®. It involves the following steps. MEDYAN code was edited to output the total number of linkers, motors, and branchers between any two filaments. The output was used to generate a graph structure with filaments as nodes and the total number of linker, motor, and brancher contacts as the edge weight.

Optimization procedure based on Blondel et al. [19] and involves two phases. A graph structure each actin filament (graph node) is assigned to its own community. Thus, we start with as many communities as there are nodes in the network. We initiate optimization moves by randomly picking a node i belonging to community $c_i$. We consider neighbors {j} of node i and calculate the change in modularity when node i is assigned to each of the communities of neighbors {j}. Node i is placed in the community where the modularity gain is maximized. This procedure is repeated till no more increase in modularity can be achieved. This marks the end of the first phase. Once completed, in the second phase, we use the community architecture at the optimal condition to generate a reduced Adjacency matrix which can be optimized further. In this graph structure, the communities from the previous iteration are defined as vertices of the graph. Edge weight between any two new vertices is defined as the sum of contacts between filaments that are part of the two communities. Connections within the same vertex lead to self-loops. By iteratively repeating phases one and two, we reduce the number of metacommunities detected. This procedure is repeated four times to understand the contact-based organization of actin filaments. Beyond four passes, we see negligible changes to the community organization of the networks studied.

**Supplementary References**


1. Popov K, Komianos J, Papoian GA. MEDYAN : Mechanochemical Simulations of Contraction and Polarity Alignment in Actomyosin Networks. PLoS Comput Biol. 2016;12: e1004877. doi:10.1371/journal.pcbi.1004877
2. Fujiwara I, Vavylonis D, Pollard TD. Polymerization kinetics of ADP- and ADP-Pi-actin determined by fluorescence microscopy. Proc Natl Acad Sci. 2007;104: 8827–8832. doi:10.1073/pnas.0702510104
3. Wachsstock DH, Schwartz WH, Pollard TD. Affinity of alpha-actinin for actin determines the structure and mechanical properties of actin filament gels. Biophys J. 1993;65: 205–14. doi:10.1016/S0006-3495(93)81059-2
4. Kovács M, Wang F, Hu A, Zhang Y, Sellers JR. Functional divergence of human cytoplasmic myosin II. Kinetic characterization of the non-muscle IIA isoform. J Biol Chem. 2003;278: 38132–38140. doi:10.1074/jbc.M305453200
5. Mahaffy RE, Pollard TD. Kinetics of the formation and dissociation of actin filament branches mediated by Arp2/3 complex. Biophys J. Elsevier; 2006;91: 3519–3528. doi:10.1529/biophysj.106.080937

6. Hansen SD, Mullins RD. VASP is a processive actin polymerase that requires monomeric actin for barbed end association. J Cell Biol. 2010;191: 571–584. doi:10.1083/jcb.201003014

7. Murphy CT, Rock RS, Spudich J a. A myosin II mutation uncouples ATPase activity from motility and shortens step size. Nat Cell Biol. 2001;3: 311–315. doi:10.1038/35060110

8. Vilfan A, Duke T. Instabilities in the transient response of muscle. Biophys J. 2003;85: 818–827. doi:10.1016/S0006-3495(03)74522-6

9. Billington N, Wang A, Mao J, Adelstein RS, Sellers JR. Characterization of three full-length human nonmuscle myosin II paralogs. J Biol Chem. Â© 2013 ASBMB. Currently published by Elsevier Inc; originally published by American Society for Biochemistry and Molecular Biology.; 2013;288: 33398–33410. doi:10.1074/jbc.M113.499848

10. Niederman R, Pollard TD. Human Platelet Myosin II . In Vitro Assembly and Structure of Myosin Filaments Formation of Platelet Myosin and Myosin Rod Filaments Determination of Platelet Myosin Solubility. J Cell Biol. 1975;67: 72–92.

11. Erdmann T, Albert PJ, Schwarz US. Stochastic dynamics of small ensembles of non-processive molecular motors: The parallel cluster model. J Chem Phys. 2013;139. doi:10.1063/1.4827497

12. Ferrer JM, Lee H, Chen J, Pelz B, Nakamura F, Kamm RD, et al. Measuring molecular rupture forces between single actin filaments and actin-binding proteins. Proc Natl Acad Sci U S A. 2008;105: 9221–6. doi:10.1073/pnas.0706124105

13. Footer MJ, Kerssemakers JWJ, Theriot JA, Dogterom M. Direct measurement of force generation by actin filament polymerization using an optical trap. Proc Natl Acad Sci. National Acad Sciences; 2007;104: 2181–2186.

14. Didonna BA, Levine AJ. Unfolding cross-linkers as rheology regulators in F-actin networks. Phys Rev E - Stat Nonlinear, Soft Matter Phys. 2007;75: 1–10. doi:10.1103/PhysRevE.75.041909

15. Li X, Ni Q, He X, Kong J, Lim S-M, Papoian GA, et al. Tensile force-induced cytoskeletal remodeling: Mechanics before chemistry. PLOS Comput Biol. 2020;16: e1007693. doi:10.1371/journal.pcbi.1007693

16. Clarke A, McQueen PG, Fang HY, Kannan R, Wang V, McCreedy E, et al. Dynamic morphogenesis of a pioneer axon in Drosophila and its regulation by Abl tyrosine kinase. Mol Biol Cell. 2020;31: 452–465. doi:10.1091/mbc.E19-10-0563

17. Clarke A, McQueen PG, Fang HY, Kannan R, Wang V, McCreedy E, et al. Abl signaling directs growth of a pioneer axon in Drosophila by shaping the intrinsic fluctuations of actin. Mol Biol Cell. 2020;31: 466–477. doi:10.1091/mbc.E19-10-0564

18. Winkelman JD, Bilancia CG, Peifer M, Kovar DR. Ena/VASP Enabled is a highly processive actin polymerase tailored to self-assemble parallel-bundled F-actin networks with Fascin. Proc Natl Acad Sci. 2014;111: 4121–4126. doi:10.1073/pnas.1322093111

19. Blondel VD, Guillaume JL, Lambiotte R, Lefebvre E. Fast unfolding of communities in large networks. J Stat Mech Theory Exp. 2008;2008. doi:10.1088/1742-5468/2008/10/P10008
